# Surveying the global landscape of post-transcriptional regulators

**DOI:** 10.1101/2021.08.09.455688

**Authors:** Kendra Reynaud, Anna McGeachy, David Noble, Zuriah Meacham, Nicholas Ingolia

## Abstract

Numerous proteins regulate gene expression by modulating mRNA translation and decay. In order to uncover the full scope of these post-transcriptional regulators, we conducted an unbiased survey that quantifies regulatory activity across the budding yeast proteome and delineates the protein domains responsible for these effects. Our approach couples a tethered function assay with quantitative single-cell fluorescence measurements to analyze ∼50,000 protein fragments and determine their effects on a tethered mRNA. We characterize hundreds of strong regulators, which are enriched for canonical and unconventional mRNA-binding proteins. Regulatory activity typically maps outside the RNA-binding domains themselves, highlighting a modular architecture that separates mRNA targeting from post-transcriptional regulation. Activity often aligns with intrinsically disordered regions that can interact with other proteins, even in core mRNA translation and degradation factors. Our results thus reveal networks of interacting proteins that control mRNA fate and illuminate the molecular basis for post-transcriptional gene regulation.

---

A dynamic network of proteins regulates the expression of messenger RNA (mRNA) to maintain homeostasis and adapt cell physiology to changing environments ^1^. This network entails a complex interplay between cis-acting sequence element encoded in the mRNA and trans-acting factors that bind the transcript to regulate its fate ^2^. RNA-binding proteins (RBPs) determine whether a given mRNA is translationally activated or repressed, localized to a specific region or compartment within the cell, or degraded ^1^. In addition, RBPs can remodel RNA structure to make it more accessible to other RBPs or enzymes and can act as chaperones to prevent RNA aggregation and misfolding ^3,4^. Thus, determining which RBPs mediate distinct regulatory programs is critical to understanding post-transcriptional control of gene expression.

Efforts to understand the diversity of RBPs and identify their mRNA targets have revealed general principles of protein-RNA interactions. Several recurring RNA binding domains (RBDs) can individually recognize 4-9 nucleotide motifs in RNA and often appear in various combinations to achieve greater specificity ^5–7^. Furthermore, protein-RNA crosslinking reveals a diverse mRNA interactome that includes many proteins without canonical RBDs ^8^, including approximately 700 high-confidence RNA-protein interactions in budding yeast ^8,9^. Reciprocally, crosslinking and immunoprecipitation (CLIP) experiments have defined the mRNA targets for hundreds of these RNA-binding proteins ^2,10,11^. Taken together, these approaches expose a dense web of interactions suggesting complex patterns of post-transcriptional regulation.

Though these high-throughput experiments have revealed principles and targets of RNA binding, we lack an understanding of the functional impact of these proteins on their target mRNAs. Traditional approaches rely on measuring how individual RBPs regulate their direct mRNA targets ^12^ and examining how these targets change when the protein is perturbed ^13^. These approaches are valuable, but they do not provide a scalable approach to characterize the regulatory networks underlying post-transcriptional regulation.

The modular architecture of regulatory RBPs ^5^ spurred the development of the tethered function assay to bypass the endogenous RNA specificity of RBPs and instead recruit them to a heterologous reporter ^14^. This approach can interrogate the regulatory effects of proteins or isolated domains, and it can reveal the function of regulatory cofactors that do not themselves bind RNA directly ^15^. In the tethering assay, candidate regulatory proteins are targeted to a reporter transcript using heterologous, bacteriophage-derived protein-RNA interaction pairs. These targeting systems utilize a specific, high-affinity interaction between a bacteriophage coat protein and a cognate RNA hairpin to recruit potential regulators to the reporter, obviating whatever direct RNA binding or indirect interactions might recruit a protein to its endogenous targets. This independence from endogenous target mRNAs and compatibility with robust reporters such as luciferase makes the tethering assay well-suited to high-throughput characterization of post-transcriptional regulators. Indeed, tethering assays have revealed over 50 regulatory proteins in a systematic analysis of 700 full-length human RBPs ^16^. It is also amenable to unbiased screening, as demonstrated by identification of almost 300 post-transcriptional regulators in trypanosomes ^17^. These studies highlight the value of tethered function assays for systematic and comprehensive study of post-transcriptional regulation.

In this study, we adapt the tethering assay to survey regulatory activity across the entire yeast proteome in an unbiased manner. Our approach allowed us to identify hundreds of proteins that modulate mRNA translation and stability, including highly active, non-canonical RBPs. We subdivided proteins and mapped their regulatory activity to particular domains and regions, in some cases uncovering effects that are not apparent in the context of the full-length protein. This fine resolution allowed us to identify Pfam protein domains and short peptide motifs that were enriched amongst the most active post-transcriptional regulators. Notably, although canonical RBPs were strongly enriched among the active regulators, the regulatory activity generally mapped outside the RBD. Taken together, our work provides a systematic, functional characterization of post-transcriptional regulators in budding yeast, thereby expanding our understanding of the complex network of proteins that control RNA metabolism.

## RESULTS

### Functional characterization of post-transcriptional regulators with a quantitative fluorescence tethering assay

We set out to functionally assess the RNA regulatory activity of proteins across the entire yeast proteome through the tethered-function assay. These tethering assays have typically used luciferase reporters ^15^, which have been adapted to measure hundreds of RBP tethering constructs in an arrayed format ^16^. In order to perform an unbiased survey of the yeast proteome not restricted to annotated RBPs, we needed to analyze libraries on a substantially larger scale.

Flow cytometry provides high-throughput, single-cell phenotypic measurements and enables large, pooled screens using fluorescence-activated cell sorting (FACS). FACS analysis of tethered function assays relies on fluorescent protein reporters ^17^, and so we devised a budding yeast tethering assay coupled to a ratiometric fluorescence readout. We target a transcript encoding a yellow fluorescent protein (YFP) by tethering it to a candidate regulatory protein, which will change YFP expression and thus yellow fluorescence. Our fluorescent reporter transcripts contained five boxB RNA hairpins in their 3′ UTR, which are recognized by the λN coat protein ^18^. In order to control for non-specific changes in cell size and physiology, we normalize these yellow fluorescence measurements against red fluorescence expressed from a control red fluorescent protein (RFP) transcript that is not targeted by λN. Changes in the ratio of fluorescence intensity between the yellow reporter and the red control provide a high-precision, quantitative measurement of specific regulatory activity affecting the targeted mRNA while controlling for global effects that also affect the non-targeted mRNA (Fig. 1a). To further control for the possibility that binding of λN itself affects the reporter, we normalize the fluorescence ratio of tethered fusion constructs against a tethered Halo protein, which exhibits no inherent regulatory effect.

**Fig. 1:**
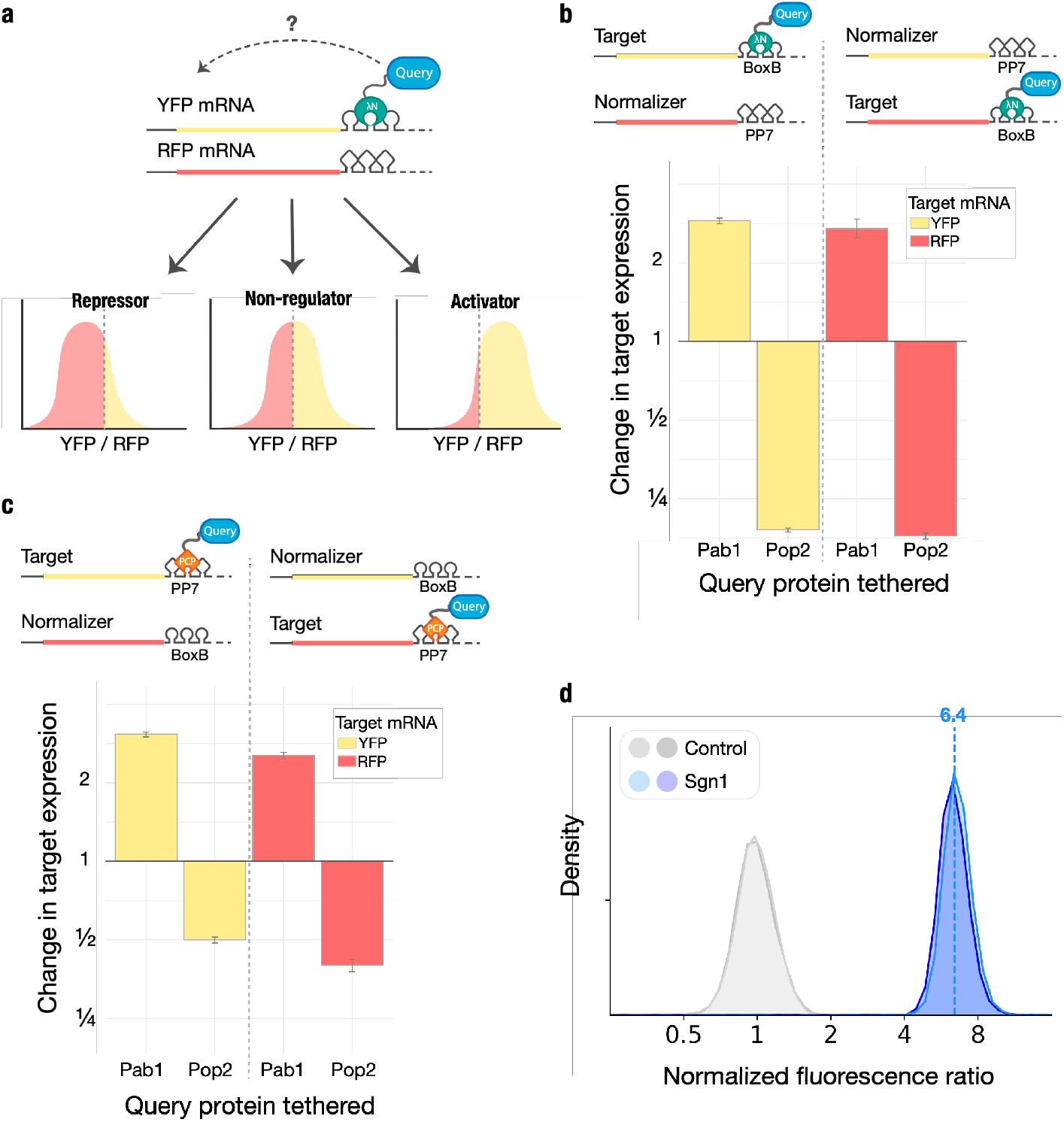
The dual reporter tethering assay reports reproducible and quantitative regulatory effects. **a**, Schematic of tethering assay with YFP reporter and RFP control (top) with expected fluorescence levels based on activity of tethered query protein (bottom). **b**, Median fluorescence ratio of Pab1 and Pop2 tethered to fluorescent protein reporter mRNAs containing boxB RNA hairpins relative to tethering an inactive Halo control (n = 3, error bars represent standard deviation). **c**, As in **b**, for Pab1 and Pop2 tethered with a single PP7 RNA hairpin (n = 3, error bars represent standard deviation). **d**, Distribution of fluorescence ratios reporting on the activity of Sgn1 in the tethering assay in two replicate samples. Dotted lines represent median YFP expression.

We validated our assay by measuring how well-characterized regulators affected reporter expression. Tethered poly(A)-binding protein (Pab1 in budding yeast) is known to enhance reporter expression by stabilizing mRNA ^14^ and promoting its translation ^19^. Conversely, the CCR4-NOT complex is responsible for the majority of cytosolic mRNA deadenylation ^20^, and tethering of the CAF1 deadenylase (Pop2 in budding yeast) greatly destabilizes target mRNAs ^21^. We observed ∼3-fold target RNA activation by tethered Pab1-λN, and ∼5-fold reporter repression by tethered Pop2-λN, relative to an inactive HaloTag-λN control (Fig. 1b). We observed quantitatively consistent effects when we switched our tethering assay to target the RFP transcript and normalized against a non-targeted YPF control, ensuring that the regulatory effects we measured were independent of our choice of fluorescent reporter protein (Fig. 1b). We further tested that they did not depend on the particular choice of the λN•boxB interaction pair by tethering Pab1 and Pop2 to reporters containing one PP7 hairpin using fusions with the PP7 coat protein (PP7cp) ^18^. Both PP7cp fusions showed similar activity on their cognate targets as λN, although Pop2-PP7cp repression appeared weaker than Pop2-λN repression, potentially due to the use of only a single PP7 hairpin (Fig. 1c). We went on to measure the activity of the RNA-binding protein Sgn1, a poorly-characterized protein that seems to be involved in regulation of translation based on co-immunoprecipitation with Pab1 and negative genetic interactions with mutations in the Pab1-interacting translation initiation factor eIF4G (Extended Data Fig. 1b) ^22^. We found that Sgn1 served as a powerful activator that upregulated YFP expression by over 6-fold relative to RFP (Fig. 1d), in addition to modestly increasing RFP levels and cell size (Extended Data Fig. 1c-f). Sgn1 tethering increased target RNA abundance roughly 2.5-fold (Extended Data Fig. 1g), and based on the larger change we see in YFP fluorescence, we infer that it activates translation as well. Taken together, these results confirm that our tethering assay provides robust and quantitative measurements of mRNA-specific regulatory activity, even in the face of additional non-specific effects on the cell, and thus provides a powerful tool for a high-throughput, proteome-wide survey of mRNA regulators.

### An unbiased survey of the yeast proteome identifies hundreds of active post-transcriptional regulators

We set out to survey the yeast proteome comprehensively for post-transcriptional regulatory activity using our two-color fluorescent tethering assay in conjunction with cell sorting and sequencing. We sought first to create a large library of λN fusion constructs and introduce this library into yeast in order to establish a pool of cells that each expressed one tethering fusion protein. FACS then allowed us to sort these cells into sub-populations according to their fluorescence phenotypes. Deep sequencing quantified the tethering constructs in each of these sorted groups. Tethering protein fusions with regulatory activity would alter the fluorescence phenotype of the host cell, shifting it into a subpopulation with an unusually low or high fluorescence ratio (Fig. 1a), and altering the distribution of the active tethering fusion across the sorted cells.

We began by generating a proteome-scale library of λN fusions that would enable unbiased discovery of regulatory proteins and identification of functional domains within these regulators. We reasoned that we could construct an unbiased λN fusion library directly from randomly fragmented genomic DNA, as budding yeast has a compact and intron-poor genome. Such a library should provide uniform representation of all proteins encoded in the proteome, based on their equal DNA copy number. Only a fraction of random genomic DNA fragments will capture the correct strand and frame of a gene, however, and so we required an additional selection for in-frame fragments. We first generated fragments by transposon-mediated tagmentation ^23,24^ and selected fragments of roughly 500 base pairs in order to capture whole protein domains, which have a typical size of ∼100 amino acids ^25^ (Fig. 2a and Extended Data Fig. 2a). We then captured these fragments by gap repair into a vector that depended on in-frame translation through the intervening sequence in order to express a downstream selectable marker (Fig. 2b). After selection, we analyzed ten individual clones and found that all ten encoded in-frame fusions, indicating that we successfully enriched for productive fusions (Extended Dataset 1). We then transferred our fragment library into a λN fusion expression vector and added random, 25-nucleotide barcodes that identify each fragment uniquely (Extended Dataset 2) ^26,27^. The mean fragment size in our barcoded λN fusion library was about 500 base pairs, consistent with the fragment size of the genomic DNA input (Fig. 2c), and we captured at least one representative fragment from roughly half of all yeast genes.

**Fig. 2:**
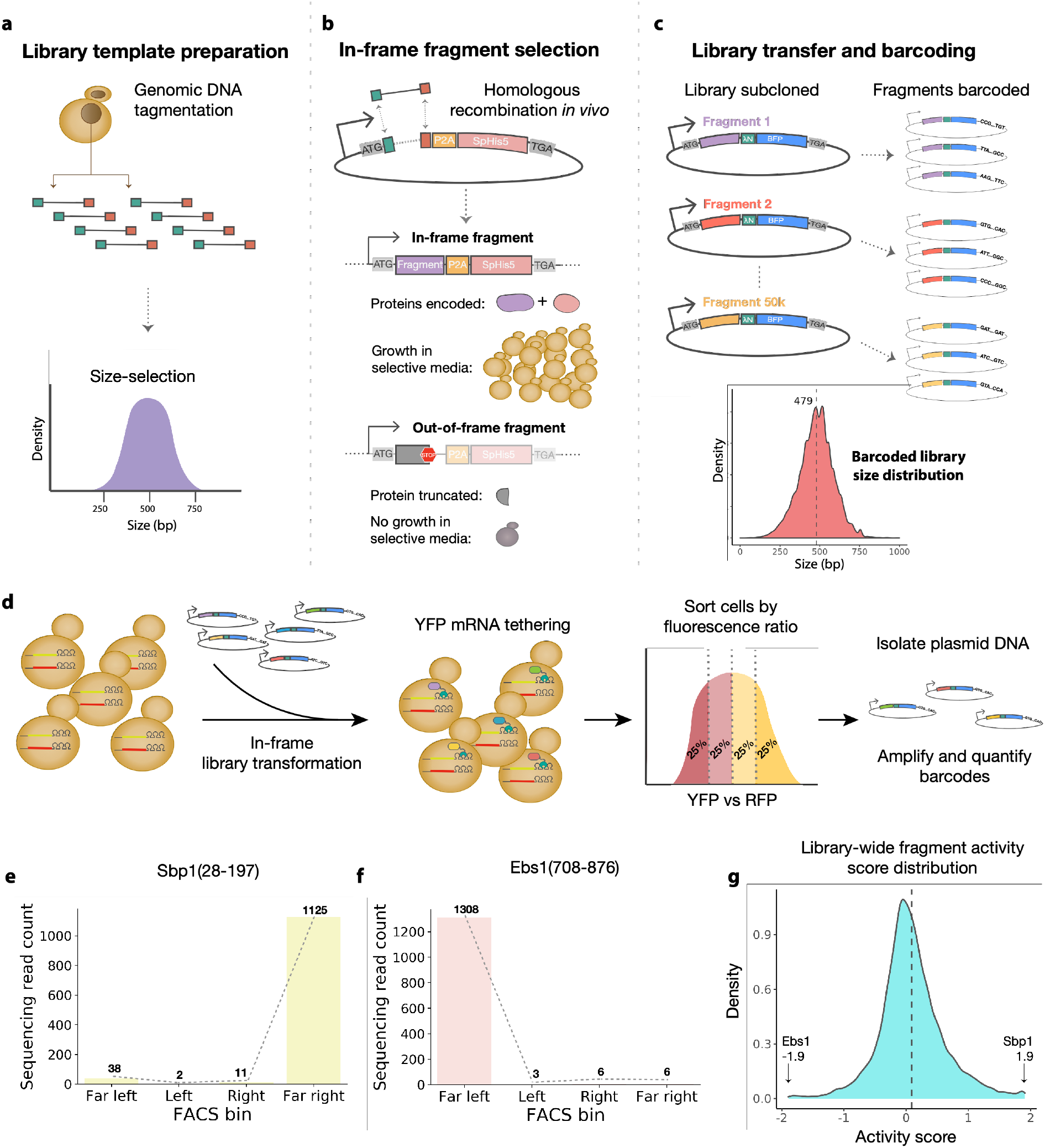
Generating an unbiased, proteome-wide survey of tethered in-frame protein fragments. **a**, S288C genomic DNA was fragmented by transposon-mediated tagmentation and selected to recover fragments with an average size of 500 base pairs. **b**, DNA fragments were cloned by *in vivo* gap repair into a plasmid containing a downstream selectable marker. Fragments containing an open reading frame in the correct phase will express a functional *S. pombe* HIS5 gene and support growth on selective media, whereas cells harboring out-of-frame fragments will fail to grow. **c**, Selected fragments were subcloned into the tethering vector with the λN and BFP proteins encoded downstream. Fragments were assigned on average three barcodes each. The barcoded library size distribution did not significantly change from in the initial input fragment library. **d**, The library was transformed into the dual-reporter yeast strain where the fragments were tethered to YFP mRNA. Phenotypic changes were captured by FACS sorting based on YFP versus RFP expression. Plasmids encoding the tethered fragments were isolated, and the barcodes associated with each fragment were amplified and then quantified through next-generation sequencing. **e**, Distribution of read counts per FACS bin for Sbp1(28-197) activator fragment. **f**, As in **e**, for Ebs1(708-876) repressor fragment. **g**, Kernel density estimate (KDE) of the library-wide activity score distribution.

We then analyzed the regulatory activity of each individual protein fragment in our library by pooled transformation, flow sorting, and sequencing. We transformed the λN fusion library into a yeast strain expressing our dual fluorescent protein reporters and sorted these transformed cells into four sub-populations of equal size according to YFP/RFP fluorescence ratio. We then isolated library plasmid DNA from our sorted cells and quantified the barcodes by next-generation sequencing (Fig. 2d). We expected activators to be enriched in bins with higher YFP/RFP ratios, while repressors should be enriched in bins with lower ratios.

Indeed, we saw certain tethering constructs that displayed a dramatic skew in their abundance across the sorted cells. For example, one particular fragment of the RNA-binding protein Sbp1 was sorted almost entirely into the highest YFP gate, indicating it strongly activated reporter expression (Fig. 2e). Conversely, a fragment of the nonsense-mediated decay factor Ebs1 acted as a strong repressor that was found almost exclusively in the lowest YFP sub-population (Fig. 2f). To quantify this enrichment, we derived an “activity score” for each fragment in our screen by computing a maximum likelihood estimate of its average fluorescence value, expressed as a *z*-score relative to the overall population. In our sorting scheme, these scores ranged from -1.9 for strong repressors like Ebs1 to +1.9 for strong activators like Sbp1. As anticipated, most fragments in our library had activity scores close to zero, indicating little or no effect on reporter transcript expression (Fig. 2g and Extended Dataset 3). Pooled screening identified active fragments from many well-known regulatory proteins such as the DEAD-box RNA helicase Ded1, a translation initiation factor ^28–30^, and Ngr1, which induces the decay of *POR1* mRNA ^31^. Our unbiased approach also uncovered novel post-transcriptional regulation in proteins with other well-characterized cellular functions, including the small heat shock chaperone Hsp26, which also has previously-identified mRNA binding activity ^32^. Furthermore, we uncovered regulatory regions in proteins of unknown function, like Her1, which may interact with ribosomes based on co-purification experiments ^33^. These results illustrate the power of our approach to discover proteins that control mRNA stability and translational efficiency and quantify how this affects gene expression.

### Full-length proteins display qualitatively similar regulatory activity as truncated fragments

We selected twelve fragments across a range of activity scores and biological functions (Fig. 3a) and tested them individually in the tethering assay by direct fluorescence measurements. All twelve fragments shifted the fluorescence ratio in the direction matching their effect in the large-scale survey (Fig. 3b), and we found a strong correlation between the magnitude of the change in YFP expression in the tethering assay and the activity score in the screen (*r =* 0.91) (Fig. 3c). This strong quantitative agreement demonstrates that the activity score derived from sorting and sequencing is an accurate measure of the regulatory effect of a fragment and can be used to discover proteins involved in post-transcriptional regulation.

**Fig. 3:**
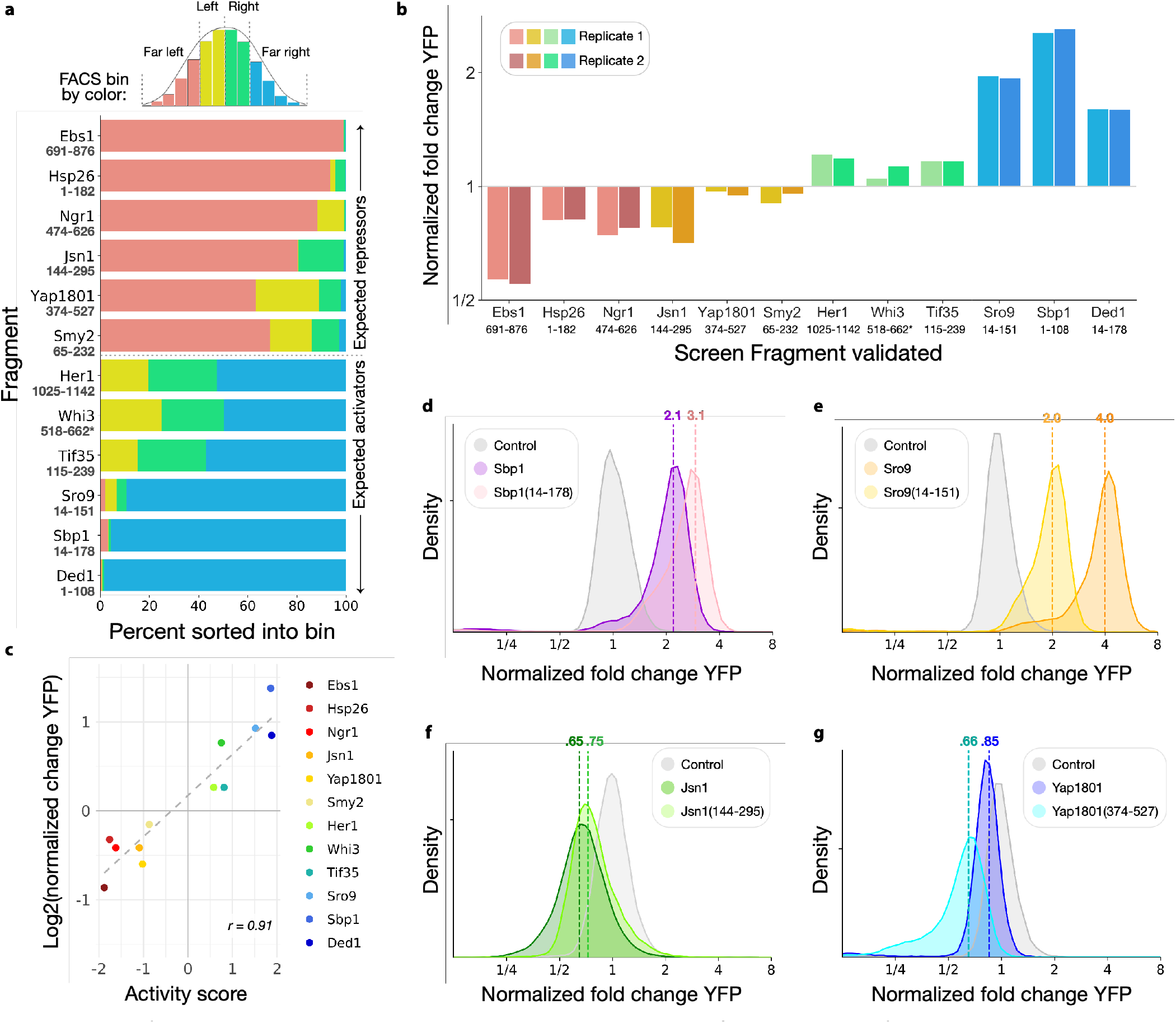
Protein fragment activity in the tethering screen represents real, verifiable regulatory function. **a**, Distribution of sequencing reads across sub-populations separated by FACS. **b**, Median activity of each protein fragment in the flow cytometry tethering assay (n = 2 per fragment). **c**, Comparison of the log_2_(difference in fluorescence ratio) and the screen activity score per fragment, *r =* 0.91. **d**, Flow cytometry measuring activity of Sbp1 and Sbp1(14-178) in the tethering assay (n = 2, one replicate per sample shown). **e**, As in **d**, for Sro9 and Sro9(14-151) (n = 2, one replicate per sample shown). **f**, As in **d**, for Jsn1 and Jsn1(144-295) (n = 2, one replicate per sample shown). **g**, As in **d**, for Yap1801 and Yap1801(374-527) (n = 2, one replicate per sample shown).

Isolated protein fragments may have different activities than the full protein from which they are derived due to the absence of regulatory domains, altered protein-protein interactions, or changes in localization, among others. We thus selected a handful of active fragments to explore how their activity relates to the wild-type protein. Full-length Sbp1 is an RBP with two RRMs in addition to an RGG-motif in the middle of the protein that recruits Pab1 ^34^. The fragment that we characterized as a ∼3-fold activator (Fig. 2e) contained only the first RRM and the RGG motif, whereas the full-length version of the protein was a weaker, ∼2-fold activator (Fig. 3d). We hypothesize that the inclusion of the second RRM interferes with its ability to recruit Pab1 as efficiently to the reporter 3′ UTR, making it a weaker activator. In other cases, the full-length protein had a stronger effect than the isolated fragment. Sro9 is an RBP that contains a La-motif and is hypothesized to activate translation through recruitment of the closed-loop-forming translation initiation complex ^35^. We identified an Sro9 fragment that activated expression ∼2-fold, whereas the full-length protein was an even more robust activator and increased reporter expression by nearly 4-fold (Fig. 3e). Tethering the entire yeast Puf-domain protein Jsn1 likewise produced a stronger repressive effect than the fragment we identified in our tethering library (Fig. 3f). In contrast, the intact version of the endocytic protein Yap1801 ^36^ was less repressive than our fragment (Fig. 3g). One possible explanation for this effect may be that Yap1801(374-527) is involved in localization of the protein to the plasma membrane where the mRNA is less actively-translated ^37^. Nonetheless, in all four cases, the full-length protein exerted an effect in the same direction as the fragment tested in our screen. Our approach is thus well suited to survey the regulatory activity contained in the native proteome and ascribe functions to RNA-binding proteins.

### Active regulators are enriched in RNA-binding proteins but not in RNA-binding domains

Our tethering assay can detect regulatory activity in truncated proteins lacking RNA-binding domains and in proteins that lack any intrinsic RNA-binding activity. This feature of the assay allows us to study the regulatory activity of co-regulators that may not bind RNA directly. Nonetheless, we did expect a substantial overlap between the post-transcriptional regulators detected in the screen and previously-known RBPs. In order to test this hypothesis, we compiled a list of budding yeast RBPs from proteins appearing in at least two of four overlapping datasets that reported RNA-protein interactions (Fig. 4a) ^38–41^. We found that fragments from these known RPBs had substantially higher absolute activity scores than the overall proteome, indicating that RBPs were more likely to show activity in our screen (Fig. 4b). This association between RBPs and regulatory activity further confirmed the relevance of our results for endogenous programs of post-transcriptional regulation controlled by these RBPs. It also raised the question whether regulatory activity was associated with the RNA-binding domains of these RBPs or with other regions of the protein.

**Fig. 4:**
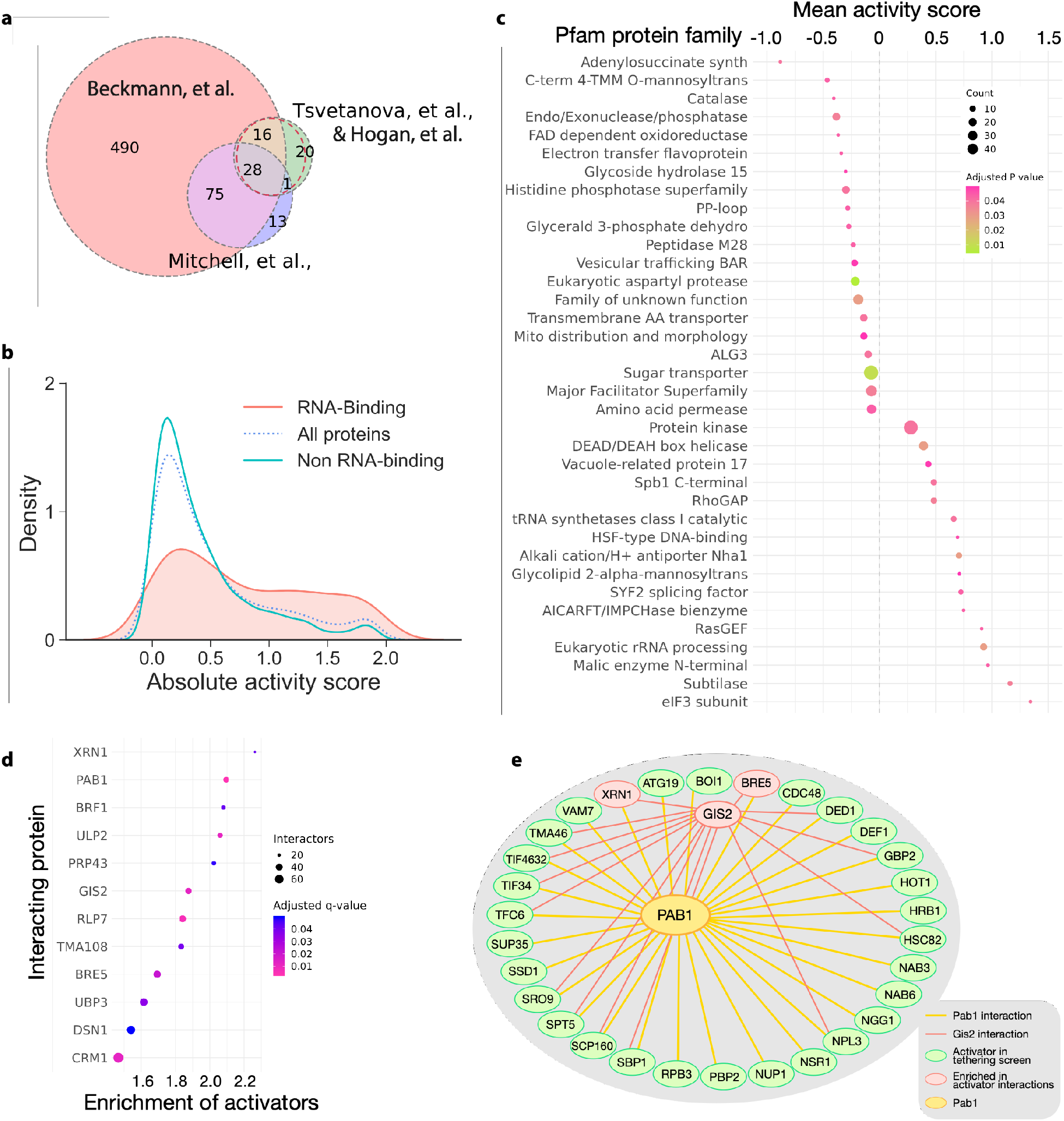
Global analyses reveal enrichment of protein domains, motifs and protein-protein interactions amongst most active screen fragments. **a**, Venn diagram of overlapping datasets identifying RBPs. **b**, KDE of absolute activity scores for RNA-binding proteins, all proteins, and non-RNA-binding proteins in the screen. **c**, Pfam protein domains significantly enriched amongst active screen fragments. **d**, Plot of proteins significantly enriched in interactions with activator fragments, x-axis represents fold-enrichment. **e**, Pab1 interactions with activator fragments (outer ring) with overlapping Gis2 interactions depicted in red.

Our tethering screen measured the activity of truncated protein fragments, and so we were able to ascribe quantitative regulatory effects to particular regions and domains of the overall protein. We were thus able to investigate which protein domains were enriched amongst the most active fragments in our screen, and in particular, whether these active regions coincided with RNA-binding domains. For each fragment, we first determined whether it contained a protein domain from the Pfam database ^42^. The boundaries of our random fragments did not align perfectly with protein domain boundaries, and so we restricted our domain enrichment analysis to library fragments containing at least 75% of some Pfam protein family. We then tested each family individually to determine whether the activity scores of fragments from that family were significantly higher or lower than the library overall (Fig. 4c).

Dozens of protein families were associated with active regulators in this analysis, and some of the strongest associations involved domains with clear connections to translation and RNA decay (Extended Dataset 6). We observed the strongest positive mean activity score amongst fragments derived from the translation initiation factor eIF3^43^. We also saw a trend for activators among the DEAD box helicase family proteins, which include translation initiation factors eIF4A and Ded1 ^44^. The endo/exonuclease/phosphatase family showed up among the strong repressors; these include certain subunits of the Ccr4-Not complex, for example ^42^. Other domains, such as Vacuole-related protein 17, which moves the vacuole along actin cables into the bud ^45^, may act through mRNA localization. We also saw many families encoding metabolic functions such as adenylosuccinate synthetase ^46^, FAD-dependent oxidoreductase, and malic enzyme N-terminal domain. Metabolic enzymes have emerged as cryptic RNA-binding proteins ^9^, and so it seems noteworthy that they appear to show regulatory activity as well. Notably, although many canonical RNA-binding domains such as RRMs appear in Pfam, they were not enriched in the active fragments. The absence of canonical RNA-recognition domains from the enriched Pfam domains indicates that RBDs are more important for mRNA target selection, and that the regions adjacent to the RNA-interacting domains typically provide regulatory activity for the RBP.

Our screen identified strong activity in fragments lacking an identifiable, folded domain. Indeed, many proteins contain intrinsically-disordered regions (IDRs) in addition to folded domains. Such IDRs play important roles in post-transcriptional regulation, and the presence of IDRs in an RBP is often an evolutionarily conserved feature of the protein ^47^. In some cases, IDRs participate in forming protein-protein interactions and can adopt a more stable conformation upon ligand binding, as in the case of the disordered N-terminus of Ded1 ^48,49^, as well as serving as flexible linkers ^50,51^. Functional IDRs can include short linear interaction motifs (SLiMs), which are often responsible for protein-protein interactions ^52^. Though SLiMs cannot be recognized in bioinformatic searches for protein domains, they may be recognizable as peptide sequence motifs.

Motivated by the possibility that SLiMs could explain regulatory effects, we searched for peptide motifs enriched in our highly-active fragments. We carried out this search using MEME, and then scanned the yeast proteome for occurrences of these motifs using FIMO ^53^. In some cases, these motifs were associated with highly repetitive elements. While these occurrences may reflect true regulatory activity, their interpretation is difficult because they are repetitive, so they were excluded. We identified six motifs that were statistically enriched amongst our activators and repressors (Extended Data Fig. 4a, b and Extended Datasets 7-8). These motifs align to genes with functions spanning many aspects of cell biology, including cell wall maintenance, cytoskeleton functions, transcription, and translation. The glutamine-rich motif (repressor motif 2 in Extended Data Fig. 4a) is particularly enriched in genes involved in mRNA metabolism, for example *NGR1, POP2*, and *PUF3*, which all have diverse roles in mRNA deadenylation and decay ^31,54,55^. Likewise, the RGG repeat in activator motif 5 (Extended Data Fig. 4b) is widespread among RBPs and linked to post-transcriptional regulation ^56^.

Regulatory RBPs often exert their effects by recruiting and activating core cellular machinery involved in translation and RNA decay. It seemed plausible that many of the active regulators we uncovered functioned in this way as well by forming protein-protein interactions with endogenous partners. We thus expected that several different active fragments from our screen might share common interactors. We intersected our library fragments with the collection of physical protein-protein interactions in the BioGRID database ^57^ and searched for proteins with a significant over-representation of activating or repressing fragments amongst their interactors. We identified a dozen such proteins (Extended Dataset 9), all enriched for interaction with activators, and most with clear ties to RNA biology (Fig. 4d). Strikingly, the poly-(A) binding protein Pab1 showed one of the highest degrees of enrichment, consistent with its central role in mRNA translation activation ^19,58^. The translation regulator Gis2 ^59,60^ was also substantially enriched in activators, and shared many interaction partners with Pab1 (Fig. 4e). Somewhat surprisingly, the exonuclease Xrn1 exhibited the strongest enrichment in interactions with activators (Fig. 4d) despite its role in mRNA decay ^61^. This enrichment may reflect a common core of mRNA-binding proteins that accompany transcripts during both translation and degradation. However, Xrn1 does promote the translation of a specific subset of transcripts encoding membrane proteins, and so this enrichment might also represent a more direct effect ^62^.

### ER/Golgi protein Gta1 is a bimodal post-transcriptional repressor

Several overlapping, C-terminal fragments of the protein Gta1 emerged as potent repressors detected in our tethering screen (Extended Dataset 5). This region of Gta1 also harbored one of the repressor-associated peptide motifs that we identified in our library (Fig. 4b and 5a). While the Gta1 protein co-purifies with the translational machinery ^33^, genetic evidence suggests that GTA1 plays a role in golgi and vesicle transport ^63,64^, and the Gta1-GFP fusion protein localizes to the ER ^65–67^. Due to its reported association with ribosomes and the presence of a repressive motif, we chose to further investigate Gta1. We generated λN fusion constructs of both Gta1(603-767), the strong repressive fragment in our library, and full-length Gta1 protein, and tested their effects on reporter expression (Fig. 5b).

**Fig. 5:**
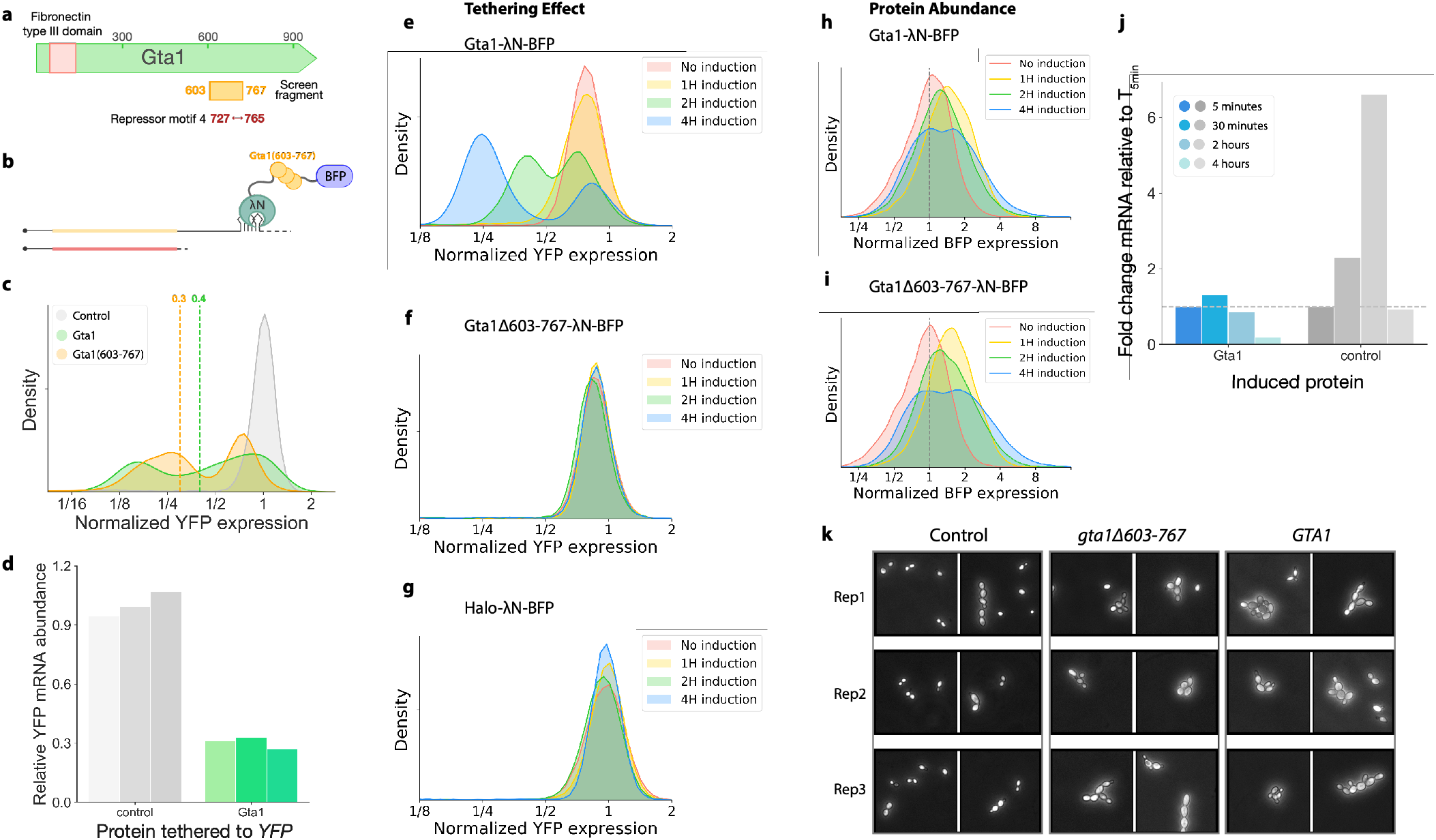
The tethering screen identifies RNA-regulatory roles of poorly-characterized proteins. **a**, Schematic representation of Gta1 protein with the C-terminal Gta1(603-767) fragment highlighted. **b**, Schematic depiction of Gta1(603-767) in the tethering assay. **c**, Flow cytometry measuring activity of Gta1 and Gta1(603-767) in the tethering assay, dotted lines represent median YFP expression (n = 2, one replicate per sample shown). **d**, RT-qPCR analysis of YFP mRNA abundance with Gta1 tethered to the 3′ UTR, normalized to a non-regulator control (n=3). **e-g**, Time course of reporter changes after induction of **e**, Gta1 **f**, Gta1Δ603-767 and **g**, Halo control tethering constructs (n=2). **h**, Change in BFP fluorescence as a measure of Gta1 expression over time, normalized to the uninduced Gta1 sample (n = 2, one replicate shown per time point). **i**, As in **h**, for Gta1Δ603-767. **j**, RT-qPCR analysis of induced *GTA1* and control mRNA over time, normalized to expression levels at 5 minutes induction. **k**, Light microscopy of yeast overexpressing *GTA1, GTA1Δ603-767*, or Halo control for 4 hours.

Both full-length Gta1 and the Gta1(603-767) fragment robustly reduced median YFP and produced a strongly bimodal distribution of reporter expression (Fig. 5c), a distinctive pattern that we did not see for any other tethering construct we examined individually. As expression of the isolated Gta1(603-767) fragment appeared to slow cell growth, we focused our analysis on full-length Gta1. We found that Gta1 tethering greatly reduced reporter mRNA abundance (Fig. 5d), suggesting that its regulatory effect principally reflected increased mRNA turnover. We next wished to track how bimodality emerged when the Gta1-λN fusion was switched on acutely, and so we expressed it from an inducible promoter. Levels of the YFP reporter began to decline within one hour of inducing the tethering construct, and clear bimodality emerged within two hours (Fig. 5e). Reporter levels in the lower peak continued to decline over the next two hours, likely reflecting the loss of pre-existing YFP through degradation or dilution. Notably, deletion of the repressor fragment that we identified in a Gta1Δ603-767-λN tethering construct abolishes this effect entirely (Fig. 5f and 5g). We thus confirm that the Gta1(603-767) region containing our repressive peptide motif is both necessary and sufficient for the regulatory effect.

We next tested whether the bimodal reporter expression we observed was a result of variation in the abundance of the Gta1 tethering fusion in the population. Indeed, we saw a broad, bimodal distribution of BFP fluorescence from the Gta1-BFP-λN construct after four hours of induction (Fig. 5h). Furthermore, it appeared that Gta1-BFP-λN levels increased more uniformly in the first hour of induction, followed by the emergence of two distinct phenotypes in this genetically homogeneous population (Fig. 5h). Notably, we saw a very similar trajectory after induction of the Gta1Δ603-767-λN tethering construct (Fig. 5i), although this tethering construct did not affect YFP expression (Fig. 5f). We also measured mRNA abundance of inducible GTA1, which quickly rose to 20 times the endogenous levels of GTA1 in an uninduced control (Extended Data Fig. 5b). Levels of the inducible mRNA did not further increase, and actually declined substantially after 4 hours in the continuous presence of the inducer (Fig. 5j). In contrast, levels of the mRNA encoding the inactive Halo-BFP-λN tethering construct used as a control in our experiments increased substantially in the two hours following induction (Fig. 5j). These mRNA abundance measurements reflect population averages, whereas flow cytometry highlights the cell-to-cell variability underlying these responses.

The heterogeneity we saw by flow cytometry led us to observe the effects of GTA1 overexpression by light microscopy. We noted that induction of full-length GTA1, or the GTA1Δ603-767 mutant lacking RNA destabilization activity, cause an atypical, elongated morphology and persistent chains and clusters of cells, akin to the filamentous or pseudohyphal growth that S. cerevisiae can undergo upon nutrient limitation (Fig. 5k) ^68,69^. Since Gta1Δ603-767 is not a strong post-transcriptional repressor, however like full-length Gta1 it appears to impact budding morphology and declines in protein abundance during prolonged transcriptional induction (Fig. 5i), the role for Gta1 in RNA turnover appears separable from its impact on budding.

### Intrinsically disordered regions mediate the activity of the translation initiation factor Ded1 and the deadenylase Ccr4

By identifying active fragments within larger proteins, we were able to distinguish the features and domains with the strongest regulatory effects. For example, our library contained many fragments of the exonuclease Ccr4, one of two proteins responsible for the deadenylase activity of the Ccr4-Not complex and thus a key factor in eukaryotic mRNA turnover ^70^. Consistent with the role of Ccr4 in deadenylation and thus mRNA decay, we identified many repressive Ccr4-derived fragments. Indeed, the median activity score of all of the assayed Ccr4 fragments was -0.5 (Fig. 6a), and the strongest repressive fragment in the screen, Ccr4(2-203), had an activity score of -1.8 (Fig. 6b). This strongly repressive fragment derived from the N-terminus of Ccr4, however, rather than the C-terminal nuclease domain ^71,72^. We obtained excellent coverage of the N-terminal half of Ccr4 (Fig. 6c), allowing us to observe that the disordered N-terminus itself, including the first 200 amino acids of the protein, yielded the strongest repressors, while the downstream folded leucine-rich repeat had little activity on its own (Fig. 6c). Our results suggest a regulatory role for the disordered N-terminus, which is not required for the nuclease activity of Ccr4 or its assembly into the Ccr4-Not complex ^72^.

**Fig. 6:**
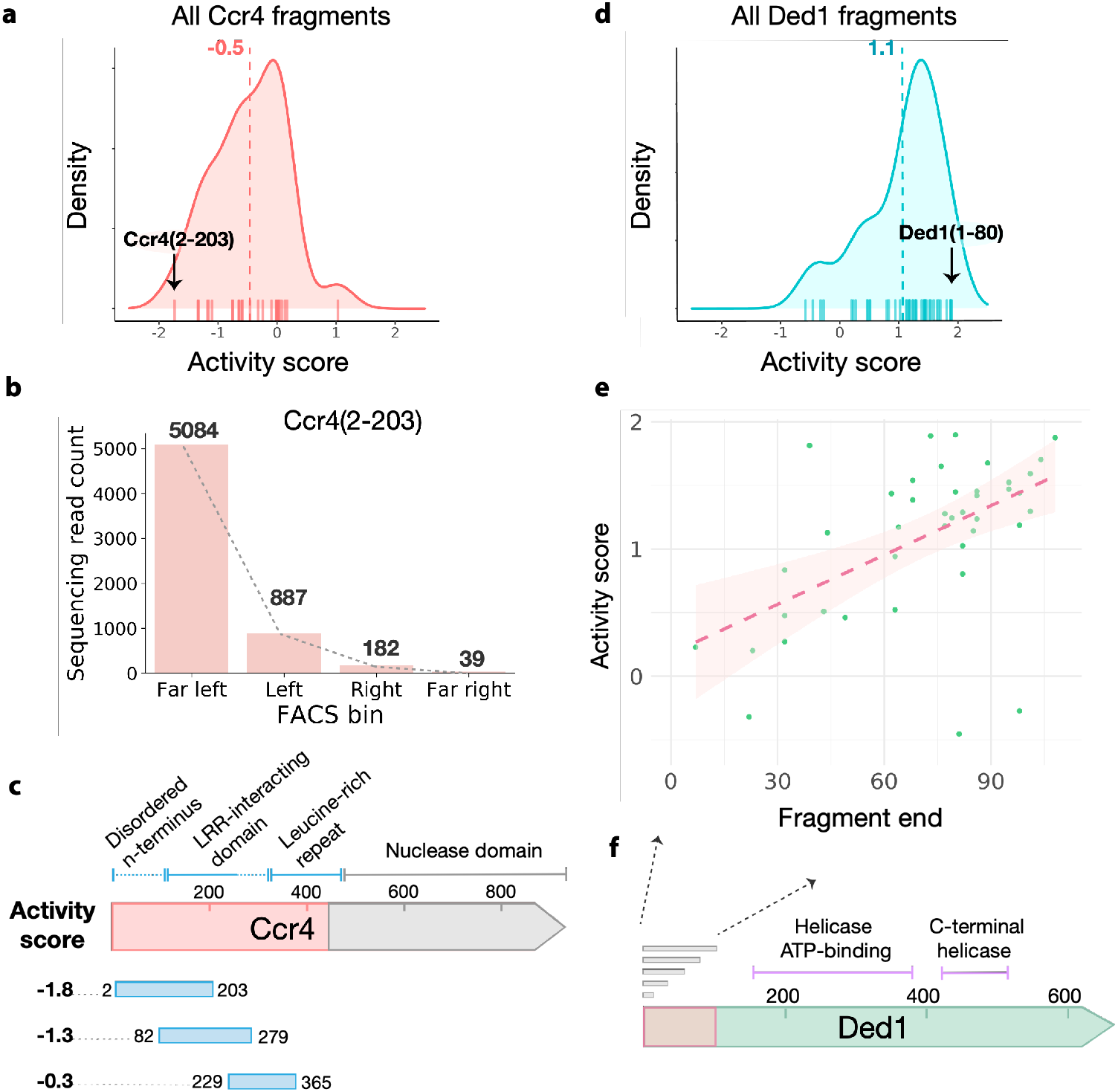
The tethering screen defines functional domain boundaries of well-characterized RBPs. **a**, KDE for all Ccr4 fragments assayed in the screen. The strongest repressor fragment, Ccr4(2-203), is indicated with the black arrow. **b**, Distribution of read counts per FACS bin for Ccr4(2-203). **c**, Schematic depiction of Ccr4 protein domains, with the strongest repressor fragment from each of the N-terminal domains shown below. **d**, As in **a**, for Ded1. The strongest activator fragment, Ded1(1-80), is indicated with the black arrow. **e**, Comparison of fragment activity score with fragment 3′ end, all fragments depicted begin at the endogenous Ded1 initiation site. **f**, Schematic depiction of Ded1 protein domains, the fragments spanning the disordered N-terminal 100 amino acid residues are indicated in **e**.

A similar pattern emerged among the regulatory fragments derived from the translation initiation factor Ded1. This protein is a highly-conserved RNA helicase of the DEAD-box family that interacts with core translation initiation factors in the cap-binding eIF4F complex and is important for translation of many yeast mRNAs ^48,73,74,49^. Ded1 fragments appeared amongst the strongest of the post-transcriptional activators in the tethering screen, consistent with its positive role in mRNA expression (Fig. 6d). We obtained excellent resolution across the disordered N-terminus of Ded1 and observed a consistent increase in activity for longer fragments (Fig. 6e and F). These results are consistent with deletion analyses that identified two distinct regions of the N-terminus required for interactions with translation initiation factors eIF4A and eIF4E ^48^. Our results indicate that full activity of Ded1 fragments in the tethered-function assay requires both of these interactions, mediated by Ded1(30-60) and Ded1(60-100), respectively. Our analysis of Ded1 and Ccr4 demonstrates the power of our approach in delineating active domains and peptide motifs within proteins and emphasizes that strong regulatory effects are often associated with disordered interaction motifs.

### The tethering screen identifies functions of the RBPs Sro9 and Cdc48

We identified and validated strong, positive post-transcriptional regulatory activity in an N-terminal fragment of the RNA-binding protein Sro9 (Fig. 3b, e). This poorly characterized protein, one of three La-motif-containing proteins in yeast, associates with translating ribosomes ^75^ and translation initiation factors ^35,76^. It appears to bind and stabilize target mRNAs enriched for functions in protein synthesis ^35^. We noted that Sro9 also contains one of the activator-associated peptide motifs we identified (Extended Data Fig. 4b, Fig. 7a), although our validated N-terminal fragment did not include this motif. In order to further dissect the regulatory activity of Sro9, we tested the full-length protein along with one truncation, Sro9(1-151), that encompassed our validated fragment, and a longer Sro9(1-251) truncation that included the activator-rich motif as well. We also tested the remaining C-terminal half of the protein, Sro9(252-434), which includes the La-motif and is implicated in RNA binding ^35^ (Fig. 7Aa). We tethered each construct to the 3′ UTR of our reporter using a λN fusion and examined their impact on its expression. Interestingly, inclusion of the activator-associated motif did not further increase the activity of the Sro9 N-terminus, although full-length Sro9 was a substantially stronger activator (Fig. 7b). The C-terminal portion alone was essentially inactive, which is consistent with our hypothesis that the RNA-binding regions of RBPs function more in cargo selection than eliciting regulatory effects. All of the fusion constructs were robustly expressed (Extended Data Fig. 7a) such that the stronger effect of the full-length protein cannot be explained by protein stability. Instead, these results point to some other way that the context of full-length Sro9 potentiates the positive effects of the N-terminal region.

**Fig. 7:**
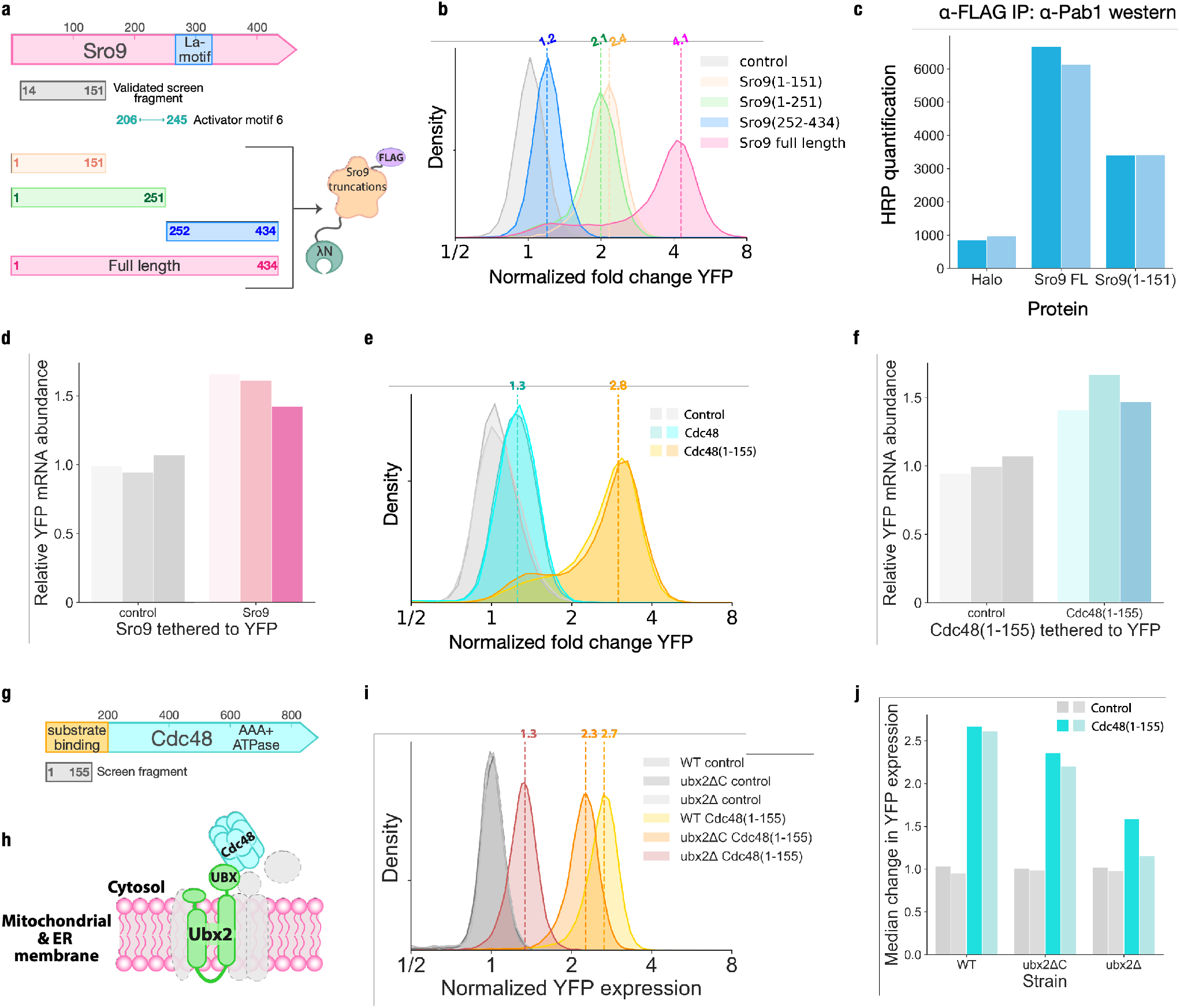
The tethering screen reveals regulatory roles of known RNA-binding proteins. **a**, Schematic representation of Sro9 protein domains and its truncations characterized in the tethering assay. **b**, Flow cytometry measuring activity of Sro9 full length and truncations in the tethering assay, dotted lines represent median YFP expression (n = 2, one replicate per sample shown). **c**, Quantification of horseradish peroxidase (HRP) western blot analysis for Pab1 enrichment in FLAG-tag immunoprecipitation eluate (n=2). **d**, RT-qPCR analysis of YFP mRNA abundance with Sro9 tethered to the 3′ UTR, normalized to a non-regulator control (n=3). **e**, As in **b**, for Cdc48 and Cdc48(1-155) (n=2). **f**, As in **d**, for Cdc48(1-155). **g**, Schematic representation of Cdc48 protein with its substrate, cofactor and ubiquitin binding N-terminal domain highlighted. **h**, Schematic depiction of Cdc48 recruitment to the ER and mitochondrial membranes through Ubx2. **i**, As in b, for Cdc48(1-155) in the wild-type, *ubx2ΔC*, and *ubx2Δ* strains (n=2, one replicate per sample shown). **j**, Quantification of median values in **i**.

Sro9 is reported to interact with several translation factors, including Pab1, which emerged as a common interaction partner for many activators in the screen (Fig. 4d, e) ^35^. We thus wanted to test whether the Sro9(1-151) fragment was sufficient for a stable Pab1 interaction. Indeed, we found that Pab1 co-purified with epitope-tagged N-terminal Sro9(1-151) (Fig. 7c and Extended Data 7b). The co-purification of Pab1 with full-length Sro9 protein was somewhat stronger, consistent with the stronger activation seen from tethered full-length Sro9 protein than from the N-terminal region alone. As Pab1 can enhance expression by stabilizing mRNAs or by promoting their translation, we wanted to test how Sro9 tethering affected mRNA abundance. We found that Sro9 increased YFP mRNA abundance by only ∼1.5-fold (Fig. 7d), indicating that increased translation explains the majority of its regulatory effect. These results are consistent with the modest quantitative changes in transcript abundance observed in *sro9Δ* yeast ^35^.

While the translation activation we observed from the Sro9 N-terminus was consistent with its known function, we also observed positive effects upon tethering proteins with little known role in mRNA regulation. Notably, we found that an N-terminal fragment of the AAA ATPase Cdc48 increased reporter expression. Cdc48 has many roles in the cell, but is linked most prominently with protein degradation, including the ER associated degradation (ERAD) pathway for quality control of transmembrane and secreted proteins ^77^. Cdc48 acts as an unfoldase that extracts proteins from membranes and complexes in order to make them available for proteasomal degradation ^78–81^. Despite these strong connections with protein turnover, Cdc48 was recently reported to interact with RNA ^9^, although its role in RNA regulation, and the identities of the specific transcripts that it binds, remain mysterious. Known functions of Cdc48 also seem to suggest that it would negatively regulate protein expression.

We were thus surprised to discover that tethering the N-terminal Cdc48(1-155) fragment to a reporter transcript robustly activated its expression (Fig. 7e and Extended Dataset 5). Furthermore, this appeared to result from enhanced translation, as reporter mRNA levels were only modestly increased by Cdc48(1-155) tethering (Fig. 7f). Interestingly, full length Cdc48 did not show this same activity (Fig. 7e). The N-terminus of Cdc48 spanning approximately the first 200 amino acids binds to substrates, cofactors, and ubiquitin, while the C-terminal domains form the hexameric AAA ATPase ^77^ (Fig. 7g). Our results thus suggest that cofactor interactions of the isolated N-terminus could increase translation. One such cofactor is the UBX-domain-containing protein Ubx2, which localizes Cdc48 to the sites of ERAD and mitochondrial protein translocation-associated degradation (mitoTAD) ^82–84^ (Fig. 7h). Deletion of the *UBX2* gene (*ubx2Δ*) reduced Cdc48(1-155) activity dramatically (Fig. 7i, j). Removal of the C-terminal UBX-domain, which mediates the interaction between Ubx2 and Cdc48 ^83^, had a smaller impact on Cdc48(1-155) activity in the tethering assay. It is possible that the localization of Cdc48(1-155) to the ER or mitochondrial surface may recruit the tethered reporter transcript and thereby modulate its expression. Alternately, the isolated N-terminus could displace binding of full-length Cdc48 from other recruitment factors on the mRNA.

## DISCUSSION

Here we report a broad and unbiased survey of the budding yeast proteome that identifies proteins controlling mRNA translation and decay. We recover a wide array of known and novel regulators, strongly enriched for RNA-binding proteins, that illustrates the breadth of post-transcriptional gene regulation. We also delineated the active regions within these proteins at domain-level resolution, which revealed that regulatory activity typically maps outside the RNA-binding domains themselves. Post-transcriptional regulators thus seem to display a modular architecture, with RNA-binding domains that determine their mRNA specificity and separate regulatory domains that modulate the expression of these target transcripts ^5,85^. We find regulatory activity associated with folded domains, but also with disordered regions, highlighting the importance of functional IDRs in post-transcriptional regulation ^85^. Indeed, the most potent repressive fragments of the Ccr4 deadenylase included its disordered N-terminus, and disordered fragments from the N-terminus of the translation initiation factor Ded1 strongly activated expression. Two broad models have emerged to explain how such IDRs might show specific molecular functions. General patterns of amino acid composition, such as interspersed acidic and aromatic residues, seem to underlie transcriptional activation by IDRs ^86–88^. Other IDRs harbor short, linear motifs (SLiMs) that act through well-defined peptide-protein docking rather than more heterogeneous interactions based on polymer properties ^89^. Both degenerate and specific peptide motifs emerged from our survey, suggesting that both modes of actions play important roles in mRNA regulation.

Many of the regulators we identified may exert their effect through their interactions with other proteins. This pattern held even in Ccr4 and Ded1, which both contain enzymatic activities that could act directly on a tethered mRNA. These interactions can affect expression of a target transcript by recruiting the large, multi-protein complexes involved in translation and mRNA decay or modulating their activity. Indeed, we observe that positive regulators were enriched for interaction with the poly-(A) binding protein Pab1, which stabilizes mRNAs and promotes their translation, suggesting that these regulators could converge onto core pathways controlling the fate of mRNAs. Similar patterns have been seen in organisms ranging from trypanosomes to humans, and so this convergence may reflect a general organizational principle of eukaryotic post-transcriptional regulation ^16,17^.

The fluorescence-based tethering assay offers a tool to further explore these regulatory networks and more broadly understand the mechanistic basis for post-transcriptional regulation. Our data also provide the key to decipher the functional consequences of the diverse RNA-binding proteome that has recently come into view.

## MATERIALS AND METHODS

### Strain construction

The dual fluorescent reporter (YFP::PP7/ RFP::boxB) strain NIY289 was constructed as follows: pNTI252 was integrated into BY4741 at URA3 to generate NIY106. pNTI476 was integrated into BY4742 at URA3 to generate NIY287. NIY106 was crossed with NIY287 to create NIY293. The dual fluorescent reporter (YFP::boxB/RFP::PP7) strain NIY293 was constructed as follows: pNTI282 was integrated into BY4741 at URA3 to generate NIY114. pNTI473 was integrated into BY4742 at URA3 to generate NIY286. NIY114 was crossed with NIY286 to create NIY293. Wild-type BY4741 was used for the in-frame library selection. The yKS090 dual reporter strain expressing the ZIF synthetic transcription factor was generated by integrating pNTI727 into the yeast XII-5 integration site ^91^. The *UBX2* mutation strains were generated as follows: We amplified the KanMX cassette from pCfB2225 with primers to generate homologous overlapping sequences to the *UBX2* locus in the yeast genome, and then integrated this cassette into one locus of *UBX2* in NIY293. We verified correct chromosomal integration by colony PCR which indicated a heterozygous *ubx2Δ/UBX2* genotype (yKS092). We then amplified the KILEU2 cassette from pCfB2189 with either homologous sequences to the remaining *UBX2* locus or to the C-terminal UBX domain of *UBX2* and then integrated these amplicons into yKS092 to create the *ubx2Δ* and *ubx2Δc* strains, respectively (yKS093 and yKS094). Genotypes were confirmed through colony PCR. Plasmids and strains are listed in Supplementary Tables 1 and 2, respectively.

### Plasmids used in this study

See Extended Data Table 1.

### Culturing conditions

Cultures for the single protein tethering assay were grown to mid-exponential growth phase at OD_600_ 0.6 then harvested via gentle centrifugation at 5000 × g for 1 minute for flow cytometry analysis with 30 minutes incubation in 4% paraformaldehyde (PFA). For in-frame fragment selection, cultures were incubated after transformation at 30 °C with shaking for 96 hours in SCD -His media with twice-daily back-dilution to avoid culture saturation. NIY293 was transformed with the in-frame tethered fragment library with the high-efficiency lithium acetate method ^93^ and then transferred to a turbidostat ^94^ for 48 hours in SCD -His media before harvesting cells. Cultures for the inducible Gta1 tethering assay were grown to stationary phase overnight, back-diluted to OD_600_ 0.1 OD and allowed to reach early exponential growth phase. The tethered proteins were then induced with 5 nM β-estradiol, then harvested and fixed in PFA as described above.

### Flow cytometric measurements and fluorescence-activated cell sorting

Expression of YFP and RFP in the tethering assay were measured using flow cytometric readout on a BD LSR Fortessa X20 with excitation by the 488mm blue laser and 561 mm yellow-green laser, captured on the FITC and PE-TexRed channels, respectively. Fluorescence measurements for 50,000 cells were collected per sample, and gates were drawn to include populations of the ∼25% cells with modal forward-and side-scatter. Fluorescence activated cell sorting was performed with an Aria Fusion sorter by gating four equal-sized populations based on the ratio of FITC and PE-TexRed emission. Approximately two million cells were sorted into each gate. The sort was performed with two technical replicate libraries from the same library transformation.

### In-frame and fragment tethering library generation

Genomic yeast DNA was tagmented using the Nextera XT DNA library prep kit from Illumina, and then size-selected with Beckman AMPureXP beads. Size selection was confirmed with an Agilent Tapestation 2200 on a High Sensitivity D1000 Screentape (Fig. S2A). BY4741 yeast were then co-transformed with the tagmented yeast gDNA and linear pKS132 and cultured as described above. After a long outgrowth in selective media, plasmids were harvested with the Zymo yeast miniprep kit. The selected fragments were then amplified by PCR with primers KS605(GTAATTATCTACTTTTTACAACAAATcctgcaggGGCTCGGAGATGTGTATAAGAGACAG) and KS630(CTGTCTCTTATACACATCTGACGCcGG AAGCGGAAGCGGAAGCCGCGCCGACGCACAAAC) designed to anneal to the Nextera XT sites introduced during tagmentation and subcloned into the SbfI-linearized tethering library vector pKS137 by Gibson Assembly. This tethering library was propagated in DH10β cells. Barcodes were then introduced by Gibson assembly of N25 randomized oligonucleotide barcodes, amplified with KS633(ACGAGGCGCGTGTAAGTTACAGGCAAGCGATCCGTCCGTAATACGACTCACTATAGCACG) and KS634(GATCCTGTAGCCCTAGACTTGATAGC CATGACTTCAACTCAAGACGCACAGATATTATAA) into the BamHI-linearized tethering library. Assembly reactions were transformed into DH10β and selected in liquid cultures at varying dilutions in order to obtain a transformant pool with roughly three barcodes per fragment. This library was transformed into NIY293 through the lithium acetate method as described in ^93^. Transformations were used to inoculate a turbidostat and grown in selective SCD -His media for 48 hours before performing FACS on live cells. Library plasmid DNA was then harvested from sorted cells with the Zymo yeast plasmid prep kit, and then barcode RNA was in-vitro transcribed with the HiScribe ™ T7 High Yield RNA kit from New England Biolabs. RNA was harvested with the previously described phenol chloroform method ^95^. All PCR reactions were performed using Q5 polymerase according to manufacturer protocols. Barcodes were amplified through a limited cycle PCR with Illumina dual-index primers. Barcodes were assigned to yeast fragment DNA with next generation sequencing using the PacBio Single Molecule Real-Time (SMRT) technology (Extended Dataset 2).

### Barcode quantification and sequencing analysis

Sequencing data was processed using Cutadapt to remove sequencing adapter sequences. HISAT2 was used to align sequencing reads to the yeast genome to identify fragment DNA. Trimmed barcodes were then counted and tabulated as described in ^96^. Barcodes that lacked at least 32 counts in one of the sorted gates were filtered out.

### RNA quantification

Total RNA was harvested from triplicate cultures of each strain using the phenol chloroform method ^95^. Quantification of *YFP* reporter RNA expression in the tethering assay was performed via RT-qPCR analysis by comparing *YFP* Ct values to *RFP* Ct values, and Ct differences for cells expressing an active tethering protein were compared to a tethered Halo protein control Ct differences. The fold change in *GTA1* expression in the induced cultures (Fig. S7A) was compared to the endogenous *GTA1* levels in the Halo-expressing control strain.

### Domain and Motif enrichment analysis

The search for domain enrichment amongst the tethered library fragments proceeded as follows: active fragments were first considered as those with an activity score of less than -1 or more than +1. Active fragments that were 90% or more similar to another fragment were considered overlapping, and the most highly sequenced from a group of overlapping fragments was used in the analysis. A given Pfam protein domain was considered represented if one or more fragments covered at least 75% of the domain. The activity scores of represented protein domains were averaged and the mean value was reported for each domain. False discovery rates (FDRs) were calculated with the Benjamini-Hochberg Procedure, and the domains with an adjusted p-value of less than 0.05 were reported in Fig. 4C.

To search for short peptide motifs enriched in our active fragments, we again considered active fragments as those with an activity score of less than -1 or greater than +1. We then ran MEME analysis to search for recurring motifs within the sequences of our active fragments ^53^. We collapsed sequences that were 50% or more similar into the same fragment in order to avoid detecting a motif multiple times within the same gene. We then used FIMO ^53^ to scan the yeast genome for occurrences of the motifs that were enriched in our active fragments. We manually removed two motifs which came from a single peptide sequence since these did not represent a consensus sequence from multiple distinct proteins, and we removed alignments that fell within highly-repetitive genomic sequences.

### Protein expression analysis via Western blotting

Total protein was isolated from mid-exponentially growing yeast by rapid capture of protein expression through 5% tricarboxylic acid treatment for ten minutes, followed by a wash in acetonitrile. The cell pellets were then dried at room temperature for 30 minutes before bead-beating in Tris-acetate-EDTA buffer for 5 minutes at room temperature. Samples were then resuspended in SDS-loading buffer from NuPage, boiled for five minutes, and loaded on 4-12% polyacrylamide Bis-Tris gels and separated by electrophoresis in MOPS buffer. Proteins were then transferred to a nitrocellulose membrane and were blocked for 1 hour in TBST with 5% bovine serum albumin. Primary antibodies were incubated with membranes at a 1-to-1000 dilution in TBST for one hour at room-temperature, washed with TBST, and then incubated for 30 minutes at room temperature with anti-rabbit and anti-mouse HRP-linked antibodies at a 1-to-10000 dilution. Membranes were developed with Pierce ECL western blotting substrate and imaged on the chemiluminescence channel on a ProteinSimple.

### Microscopy

Mid-exponential phase cells were harvested by gentle centrifugation and then fixed for 30 minutes in 4% PFA. The cells were washed in 1x PBS buffer and visualized with a Leica DM IL LED microscope at 40X magnification. Fields of view for saved images were randomly selected.

## ACKNOWLEDGEMENTS

We thank Shintaro Iwasaki and Gloria Brar for insightful comments and suggestions. We thank Ryan Muller for library plasmids, Samantha Fernandez for assistance with computational data visualization, and Joe Lobel along with other members of the Ingolia lab for thoughtful scientific discussion. This work was supported by NIH grants DP2 CA195768 (N.T.I.) and R01 GM130996 (N.T.I.) and by the Rose Hills Innovator Program. This work used the Vincent J. Coates Genomics Sequencing Laboratory at UC Berkeley, supported by NIH S10 OD018174 Instrumentation Grant, as well as the Flow Cytometry Facility at UC Berkeley, the UC Berkeley DNA Sequencing Facility, and the UC Davis Proteomics Core Facility.

## AUTHOR CONTRIBUTIONS

Conceptualization: KR, AM, NI

Data curation: KR, NI

Formal analysis: KR, DN, NI

Funding acquisition: NI

Investigation: KR, AM, ZM

Methodology: KR, AM, NI

Project administration: KR, NI

Resources: NI

Supervision: NI

Validation: KR

Visualization: KR

Writing -original draft: KR, NI

Writing -review & editing: KR, AM, NI

## DECLARATION OF INTERESTS

The authors declare no competing interests.

## DATA AVAILABILITY

- High-throughput sequencing data has been deposited with the NCBI Short Read Archive under BioProject <u>PRJNA579846</u>.
- Custom software and scripts are available from Zenodo at https://doi.org/10.5281/zenodo.4963329
- Plasmids will be made available through Addgene.
- Yeast strains described in this study are available upon request from the corresponding author.

## Notes

https://www.ncbi.nlm.nih.gov/Traces/study/?acc=PRJNA579846

https://doi.org/10.5281/zenodo.4963329

### Competing Interest Statement

The authors have declared no competing interest.

