## Supplemental Figures and Tables for "Surveying the global landscape of post-transcriptional regulators"

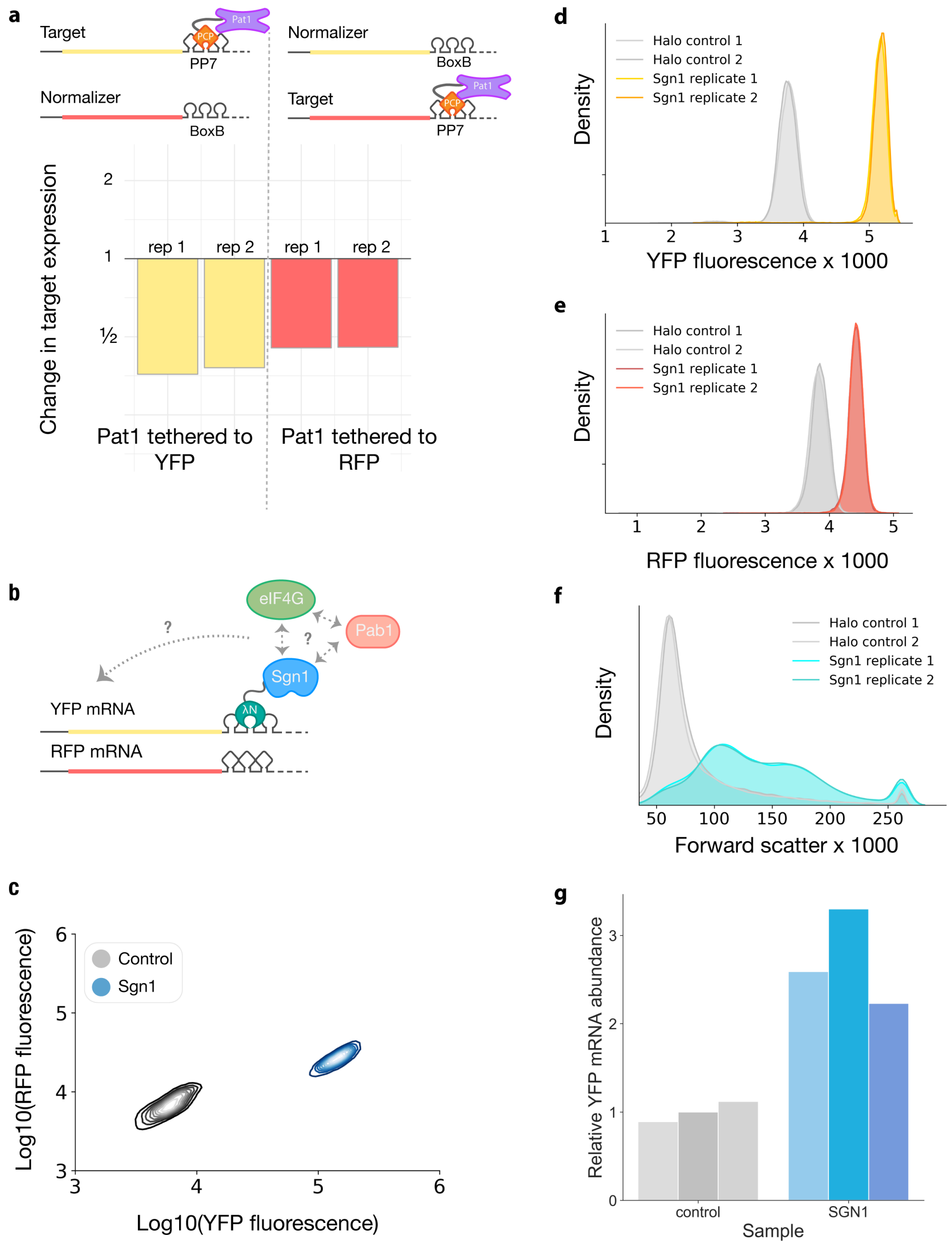


### Extended Data Figure 1

**a**, Pat1 activity tethered to 3′ UTR of both fluorescent reporters with PP7 is reproducible between replicates and fluorophores. **b**, Schematic representation of Sgn1 recruiting Pab1 and eIF4G in the tethering assay. **c**, Comparison of RFP and YFP fluorescence with Sgn1 or a non-regulator control tethered to YFP (n = 3, one representative replicate depicted. **d**, YFP fluorescence with Sgn1 and the non-regulator Halo protein tethered to the 3′ UTR. **e**, RFP fluorescence with Sgn1 and the non-regulator Halo protein tethered to the 3′ UTR of YFP. **f**, Forward scatter fluorescence of Sgn1 and control protein expressing cells indicates larger cell size in the Sgn1 samples. **g**, RT-qPCR analysis of YFP mRNA levels with Sgn1 or the control tethered to the 3′ UTR.

**a** Fragment input library size distribution:

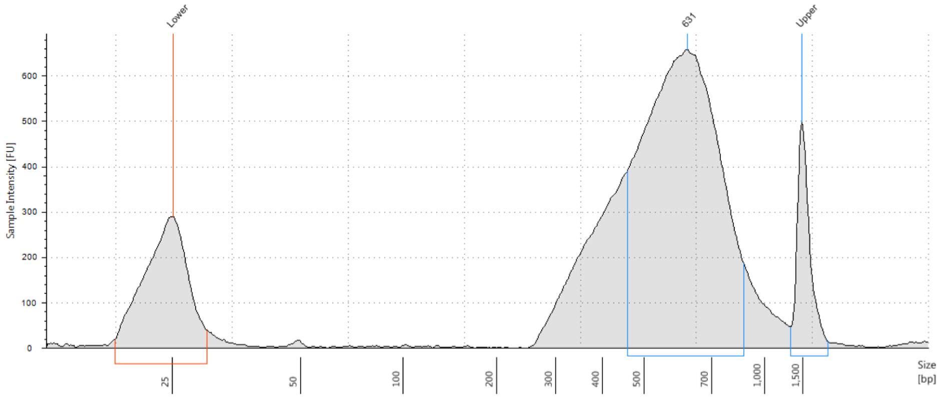


### Extended Data Figure 2

**a**, Bioanalyzer analysis of tagmented genomic DNA library size distribution.

**a**

| barcode | ORF | gene | fragment start | far left counts | left counts | right counts | far right counts | unsorted counts | activity score |
|---|---|---|---|---|---|---|---|---|---|
| GGATACTAGCGCTCTGGTCATCTTA | YDR206W | EBS1 | chr04:864649- | 1178 | 0 | 3 | 2 | 10 | -1.90 |
| TGTGTTGTGGGTAGTTATTGAATAG | YBR072W | HSP26 | chr02:382528- | 2232 | 50 | 93 | 9 | 29 | -1.71 |
| CGCCTTTATTGGGTTTATCTGTGGG | YBR212W | NGR1 | chr02:649731- | 1508 | 186 | 8 | 5 | 21 | -1.62 |
| CGGTGCGTATTTCGGCTCGGTGTGC | YJR091C | JSN1 | chr10:596252+ | 427 | 2 | 98 | 5 | 6 | -1.28 |
| GGTGCACCATCCCGGACTATCGTG | YHR161C | YAP1801 | chr08:420484+ | 2011 | 710 | 13 | 175 | 43 | -1.10 |
| GGGGTAGGTACGAATAGACCCGTGG | YBR172C | SMY2 | chr02:580677+ | 10527 | 2584 | 1697 | 420 | 294 | -1.09 |
| GCGGGATCCACTGGGCGGCGAGATA | YOR227W | HER1 | chr15:766230- | 64 | 272 | 144 | 430 | 17 | 0.63 |
| TTTCTGATTGACGCTAAGCTTACTG | YNL197C | WHI3 | chr14:267662+ | 5 | 493 | 510 | 995 | 30 | 0.82 |
| GATAACACTTGCAGACCAATCTTTT | YDR429C | TIF35 | chr04:1324585+ | 46 | 1243 | 1136 | 2637 | 66 | 0.84 |
| ACCGGGTTTTAAATTTTCTAAATCA | YCL037C | SRO9 | chr03:58019+ | 239 | 5 | 36 | 2101 | 36 | 1.49 |
| AAGTTCGGTCTGCACTAACCGTAAT | YOR204W | DED1 | chr15:723124- | 5 | 0 | 10 | 1525 | 22 | 1.90 |
| TCTTCACTTACGGTGGAGTGGAATT | YHL034C | SBP1 | chr08:33544+ | 0 | 0 | 0 | 1553 | 690 | 1.92 |

**b**

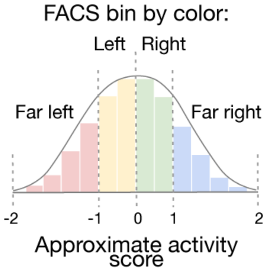


### Extended Data Figure 3

**a**, FACS bin counts and activity scores for validated screen fragments. **b**, Schematic key depicting FACS bins based on position and color.

**a**

| | Repressors: Enriched motifs | Motif E-value | Most significant gene | Gene q-value | Count |
|---|---|---|---|---|---|
| 1 | 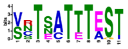 | $7.3 \times 10^{-88}$ | YKR102W, FLO10 | $3.39 \times 10^{-8}$ | 106 |
| 2 | 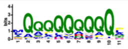 | $1.8 \times 10^{-14}$ | YFR008W, FAR7 | $4.26 \times 10^{-10}$ | 524 |
| 3 | 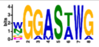 | $2.7 \times 10^{-14}$ | YML010W, SPT5 | $5.48 \times 10^{-05}$ | 1 |
| 4 | 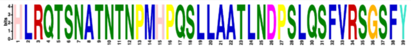 | $1.1 \times 10^{-6}$ | YEL043W, GTA1 | $2.06 \times 10^{-43}$ | 3 |
| 5 | 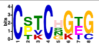 | $5.9 \times 10^{-6}$ | YFL016C, MDJ1 | $7.41 \times 10^{-05}$ | 9 |
| 6 | 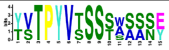 | $1.4 \times 10^{-4}$ | YKR102W, FLO10 | $9.22 \times 10^{-13}$ | 483 |

**b**

| | Activators: Enriched motifs | Motif E-value | Most significant gene | Gene q-value | Count |
|---|---|---|---|---|---|
| 1 | 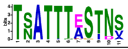 | $1.2 \times 10^{-513}$ | YKR103W, PRY2 | $1.06 \times 10^{-3}$ | 157 |
| 2 | 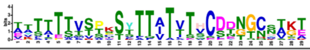 | $8.4 \times 10^{-69}$ | YOL155C, HPF1 | $2.04 \times 10^{-25}$ | 563 |
| 3 | 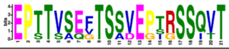 | $1.7 \times 10^{-54}$ | YJR151C, DAN4 | $1.59 \times 10^{-20}$ | 38 |
| 4 | 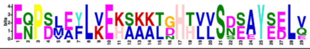 | $2.5 \times 10^{-41}$ | YDR150W, NUM1 | $1.61 \times 10^{-25}$ | 9 |
| 5 | 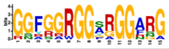 | $6.6 \times 10^{-37}$ | YDL014W, NOP1 | $2.12 \times 10^{-12}$ | 63 |
| 6 | 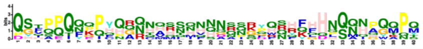 | $2.3 \times 10^{-35}$ | YCL037C, SRO9 | $5.31 \times 10^{-36}$ | 618 |

### Extended Data Figure 4

**a**, Peptide motifs significantly enriched amongst repressor screen fragments. Counts represent significant occurrences of that motif in the yeast genome. **b**, As in **a**, for the activator screen fragments.

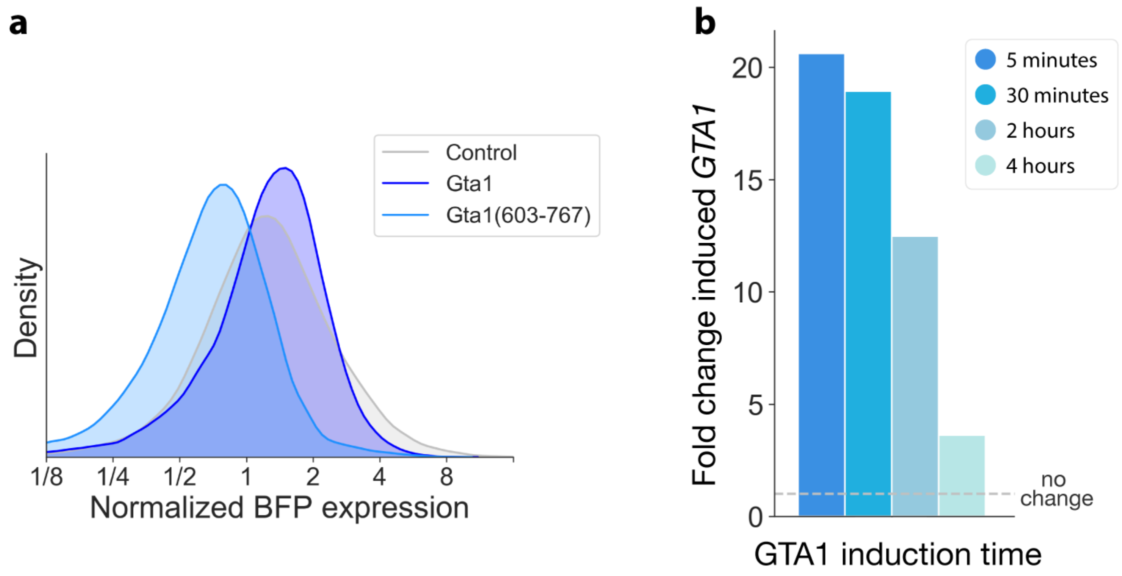


### Extended Data Figure 5

**a**, Histogram of BFP fluorescence as a measure for control, Gta1 and Gta1(603-767) expression and stability in the tethering assay, normalized to control BFP levels. Dotted line represents median BFP levels of Gta1(603-767) (n = 2, one replicate per sample shown). **b**, Fold change of induced *GTA1* over time relative to uninduced endogenous *GTA1* expression (n = 3).

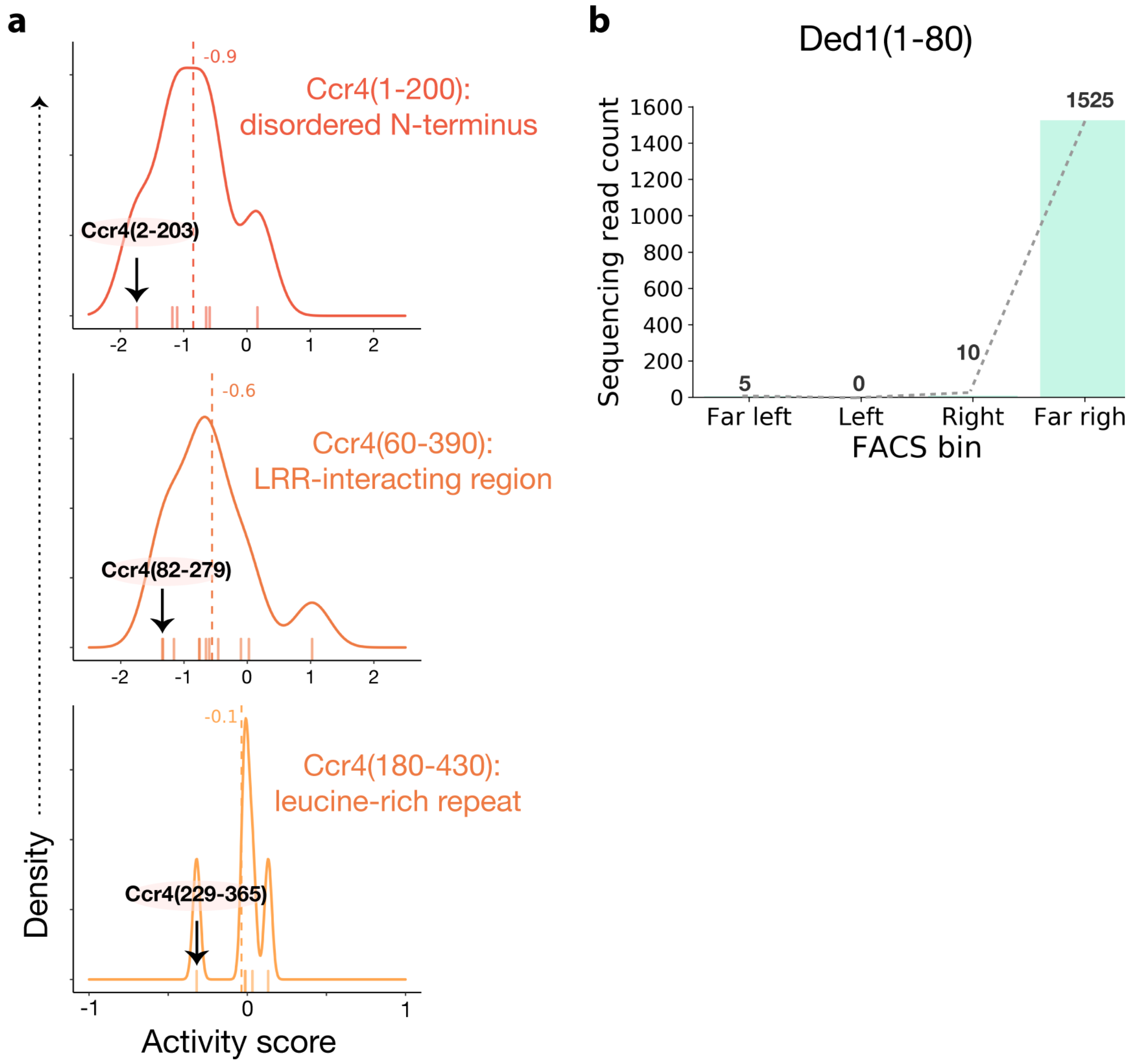


### Extended Data Figure 6

**a**, KDE for Ccr4(1-200) disordered N-terminus domain (top), Ccr4(60-390) Leucine-rich repeat interacting domain (middle), and Ccr4(180-340) leucine-rich repeat domain (bottom). **b**, As in Fig. 6b, for Ded1(1-80).

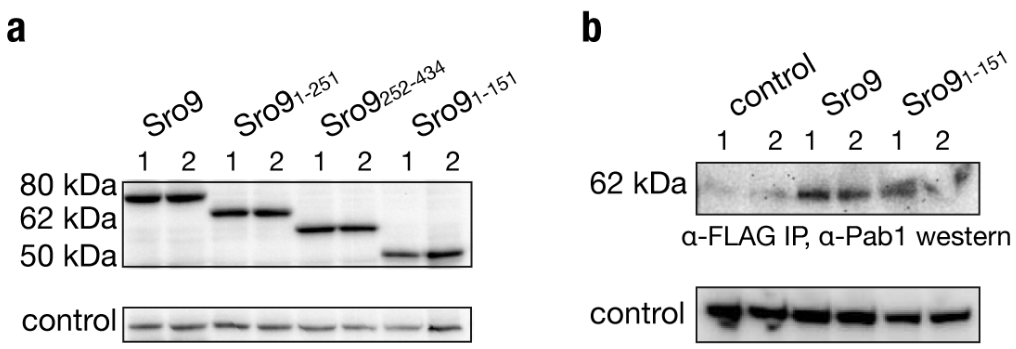


### Extended Data Figure 7

**a**, Western blot analysis of Sro9 full length and truncation protein expression in the tethering assay. **b**, Western blot analysis of Pab1 enrichment in FLAG-tag protein purification eluate.

### Extended Data Table 1: Plasmids used in this study.

| Plasmid name | Purpose | Expression cassette | Source |
|---|---|---|---|
| pKS038 | Sgn1 tethering vector | pPGK1::SGN1-λN:-SpHis5::tADH1 ARS/CEN | This study |
| pKS109 | Halo control tethering vector | pPGK1::Halo-λN-SpHis5::tADH1 ARS/CEN | This study |
| pKS132 | Library in-frame selection vector | pPGK1::λN-P2A-SpHis5::tAgTEF ARS/CEN | This study |
| pKS137 | Library tethering vector | pPGK1::λN-T2A*-1xFLAG-BFP-NES::tADH1 pAgTEF::SpHIS5::tAgTEF ARS/CEN | This study |
| pKS151 | Ded1(14-178) tethering vector | pPGK1::Ded1(14-178)-λN-T2A*-1xFLAG-BFP-NES::tADH1 pAgTEF::SpHIS5::tAgTEF ARS/CEN | This study |
| pKS152 | Ebs1(691-876) tethering vector | pPGK1::Ebs1(691-876)-λN-T2A*-1xFLAG-BFP-NES::tADH1 pAgTEF::SpHIS5::tAgTEF ARS/CEN | This study |
| pKS153 | Hsp26(1-182) tethering vector | pPGK1::Hsp26(1-182)-λN-T2A*-1xFLAG-BFP-NES::tADH1 pAgTEF::SpHIS5::tAgTEF ARS/CEN | This study |
| pKS154 | Ngr1(474-626) tethering vector | pPGK1::Ngr1(474-626)-λN-T2A*-1xFLAG-BFP-NES::tADH1 pAgTEF::SpHIS5::tAgTEF ARS/CEN | This study |
| pKS155 | Tif35(115-239) tethering vector | pPGK1::Tif35(115-239)-λN-T2A*-1xFLAG-BFP-NES::tADH1 pAgTEF::SpHIS5::tAgTEF ARS/CEN | This study |
| pKS156 | Gta1(603-767) tethering vector | pPGK1::Gta1(603-767)-λN-T2A*-1xFLAG-BFP-NES::tADH1 pAgTEF::SpHIS5::tAgTEF ARS/CEN | This study |
| pKS160 | Her1(1025-1142) tethering vector | pPGK1::Her1(1025-1142)-λN-T2A*-1xFLAG-BFP-NES::tADH1 pAgTEF::SpHIS5::tAgTEF ARS/CEN | This study |
| pKS171 | Sro9(14-151) tethering vector | pPGK1::Sro9(14-151)-λN-T2A*-1xFLAG-BFP-NES::tADH1 pAgTEF::SpHIS5::tAgTEF ARS/CEN | This study |
| pKS173 | Sbp1(1-108) tethering vector | pPGK1::Sbp1(1-108)-λN-T2A*-1xFLAG-BFP-NES::tADH1 pAgTEF::SpHIS5::tAgTEF ARS/CEN | This study |
| pKS174 | Smy2(65-232) tethering vector | pPGK1::Smy2(65-232)-λN-T2A*-1xFLAG-BFP-NES::tADH1 pAgTEF::SpHIS5::tAgTEF ARS/CEN | This study |
| pKS176 | Jsn1(144-295) tethering vector | pPGK1::Jsn1(144-295)-λN-T2A*-1xFLAG-BFP-NES::tADH1 pAgTEF::SpHIS5::tAgTEF ARS/CEN | This study |
| pKS177 | Cdc48 tethering vector | pPGK1::CDC48-λN-T2A*-1xFLAG-BFP-NES::tADH1 pAgTEF::SpHIS5::tAgTEF ARS/CEN | This study |
| pKS179 | Yap1801 tethering vector | pPGK1::YAP1801-λN-T2A*-1xFLAG-BFP-NES::tADH1 pAgTEF::SpHIS5::tAgTEF ARS/CEN | This study |

| | | | |
|---|---|---|---|
| pKS180 | Sbp1 tethering vector | pPGK1::SBP1-λN-T2A*-1xFLAG-BFP-NES::tADH1 pAgTEF::SpHIS5::tAgTEF ARS/CEN | This study |
| pKS182 | Sro9 tethering vector | pPGK1::SRO9-λN-T2A*-1xFLAG-BFP-NES::tADH1 pAgTEF::SpHIS5::tAgTEF ARS/CEN | This study |
| pKS183 | Gta1 tethering vector | pPGK1::GTA1-λN-T2A*-1xFLAG-BFP-NES::tADH1 pAgTEF::SpHIS5::tAgTEF ARS/CEN | This study |
| pKS190 | *Gta1Δ603-767* tethering vector | pPGK1::Gta1Δ603-767-λN-T2A*-1xFLAG-BFP-NES::tADH1 pAgTEF::SpHIS5::tAgTEF ARS/CEN | This study |
| pKS192 | Sro9-3xFLAG tethering vector | pPGK1::SRO9-3xFLAG-λN-T2A*-1xFLAG-BFP-NES::tADH1 pAgTEF::SpHIS5::tAgTEF ARS/CEN | This study |
| pKS193 | Sro9(1-151)-3xFLAG tethering vector | pPGK1::SRO9(1-151)-3xFLAG-λN-T2A*-1xFLAG-BFP-NES::tADH1 pAgTEF::SpHIS5::tAgTEF ARS/CEN | This study |
| pKS194 | Sro9(1-251)-3xFLAG tethering vector | pPGK1::SRO9(1-251)-3xFLAG-λN-T2A*-1xFLAG-BFP-NES::tADH1 pAgTEF::SpHIS5::tAgTEF ARS/CEN | This study |
| pKS196 | Halo-3xFLAG tethering vector | pPGK1::Halo-3xFLAG-λN-T2A*-1xFLAG-BFP-NES::tADH1 pAgTEF::SpHIS5::tAgTEF ARS/CEN | This study |
| pKS207 | Gta1 inducible vector | pGAL1::GTA1-λN-T2A*-1xFLAG-BFP-NES::tADH1 | This study |
| pKS208 | *Gta1Δ603-767* inducible vector | pGAL1::Gta1Δ603-767-λN-T2A*-1xFLAG-BFP-NES::tADH1 | This study |
| pKS232 | Halo inducible vector | pGAL1::Halo-λN-T2A*-1xFLAG-BFP-NES::tADH1 | This study |
| pNTI282 | YFP-boxB vector | pCMV::eGFP::5xboxB::poly(A) BGH | This study |
| pNTI473 | RFP-PP7 vector | pPGK1::mCherry::3xPP7::tADH1 | This study |
| pHES795 | ZIF268 synthetic transcription factor vector | Zif268 DBD-hPR LBD-MSN2 AD | [90] |
| pHES840 | pGAL1 inducible promoter vector | pGAL1::YFP | [90] |
| pCfB2189 | Leu+ vector | KILEU2 at integration site X-3 | [91] |
| pCfB2225 | Kan+ vector | KanMX at integration site XII-2 | [91] |
| pCfB2337 | Hygromycin+ vector | HphMX at integration site XII-5 | [91] |
| pNTI114 | YFP::boxB integration vector | CCR5_3'HR:P_CMV::eGFP::polyA_BGH:CCR5_5'HR | This Study |
| pNTI252 | RFP::boxB integration vector | pFA6a P(PGKI)::mCherry:boxB::T(ADH1) CaURA3 URA3int | This Study |

| | | | |
|---|---|---|---|
| pNTI473 | RFP::PP7 integration vector | pFA6a P(PGK1)::mCherry:pp7(x3):T(ADH1) CaURA3 URA3int | This Study |
| pNTI476 | YFP::PP7 vector | pFA6a P(PGK1)::YFP:pp7:T(ADH1) CaURA3 URA3int | This Study |
| pNTI729 | ZIF268 synthetic transcription factor integratable vector | pADH1::ZIF268::C.albicans tADH1 at site XII-6 | [92] |

### Extended Data Table 2: Strains used in this study.

| **Name** | **Genotype** | **Purpose** | **Source** |
|---|---|---|---|
| NIY106 | MATα his3Δ1 leu2Δ0 lys2Δ0 MET15 pPGKI::mCherry:boxB:CaUra3 | RFP::boxB haploid strain | This study |
| NIY111 | MATa his3Δ1 leu2Δ LYS2 met15Δ ura3Δ0 | BY4741 MATa WT yeast | ThermoFisher |
| NIY112 | MATα his3Δ leu2Δ0 lys2Δ0 MET15 ura3Δ0 | BY4742 MATα WT yeast | ThermoFisher |
| NIY114 | MATa his3Δ1 leu2Δ LYS2 met15Δ pPGKI::YFP:boxB:CaUra3 | YFP::boxB haploid strain | This study |
| NIY286 | MATα his3Δ1 leu2Δ0 lys2Δ0 MET15 pPGKI::mCherry:PP7:CaUra3 | RFP::PP7 haploid strain | This study |
| NIY287 | MATa his3Δ1 leu2Δ LYS2 met15Δ pPGKI::YFP:pp7:CaUra3 | YFP::PP7 haploid strain | This Study |
| NIY289 | MATα/MATa his3Δ1/his3Δ1 leu2Δ0/ leu2Δ0 lys2Δ0/LYS2 MET15/met15Δ pPGKI::mCherry:boxB:CaUra3/ pPGKI::YFP:pp7:CaUra3 | YFP::PP7/RFP::boxB dual reporter strain | This Study |
| NIY293 | MATα/MATa his3Δ1/his3Δ1 leu2Δ0/ leu2Δ0 lys2Δ0/LYS2 MET15/met15Δ pPGKI::mCherry:pp7:CaUra3/ pPGKI::YFP:boxB:CaUra3 | YFP::boxB/RFP::PP7 dual reporter strain | This study |
| yKS090 | MATα/MATa his3Δ1/his3Δ1 leu2Δ0/ leu2Δ0 lys2Δ0/LYS2 MET15/met15Δ HygR pPGKI::mCherry:pp7:CaUra3/ pPGKI::YFP:boxB:CaUra3 pADH1::ZIF268::tADH1 | Dual reporter strain NIY293 with ZIF268 synthetic transcription factor integrated at XII-5 site | This study |
| yKS092 | MATα/MATa his3Δ1/his3Δ1 LEU2/ leu2Δ0 lys2Δ0/LYS2 MET15/met15Δ *ubx2Δ*/UBX2 KanR pPGKI::mCherry:pp7:CaUra3/ pPGKI::YFP:boxB:CaUra3 | Dual reporter with 1 copy *ubx2Δ*, 1 copy UXB2 | This Study |
| yKS093 | MATα/MATa his3Δ1/his3Δ1 LEU2/ leu2Δ0 lys2Δ0/LYS2 MET15/met15Δ *ubx2Δ*/*ubx2Δ* KanR pPGKI::mCherry:pp7:CaUra3/ pPGKI::YFP:boxB:CaUra3 | Dual reporter *ubx2Δ* strain | This study |
| yKS094 | MATα/MATa his3Δ1/his3Δ1 LEU2/ leu2Δ0 lys2Δ0/LYS2 MET15/met15Δ *ubx2Δ*/*ubx2Δubx* KanR pPGKI::mCherry:pp7:CaUra3/ pPGKI::YFP:boxB:CaUra3 | Dual reporter *ubx2Δc* (no UBX domain) strain | This Study |
